# Impairments in fear extinction memory and basolateral amygdala plasticity in the TgF344-AD rat model of Alzheimer’s disease are distinct from non-pathological aging

**DOI:** 10.1101/2022.03.12.484093

**Authors:** Caesar M. Hernandez, Nateka L. Jackson, Abbi R. Hernandez, Lori L. McMahon

## Abstract

Fear-based disorders such as posttraumatic stress disorder (PTSD) steepen age-related cognitive decline and doubles the risk for developing Alzheimer’s disease. Due to the seemingly hyperactive properties of fear memories, PTSD symptoms can worsen with age. Perturbations in the synaptic circuitry supporting fear memory extinction are key neural substrates of PTSD. The basolateral amygdala (BLA) is a medial temporal lobe structure critical in the encoding, consolidation, and retrieval of fear memories. As little is known about fear extinction memory and BLA synaptic dysfunction within the context of aging and AD, the goal of this study was to investigate how fear extinction memory deficits and basal amygdaloid nucleus (BA) synaptic dysfunction differentially associate in non-pathological aging and AD in the TgF344AD (TgAD) rat model of AD. Young, middle-aged, and older-aged WT and TgAD rats were trained on a delay fear conditioning and extinction procedure prior to *ex vivo* extracellular field potential recording experiments in the BA. Relative to young WT rats, long-term extinction memory was impaired, and in general, associated with a hyperexcitable BA and impaired LTP in TgAD rats at all ages. In contrast, long-term extinction memory was impaired in aged WT rats and associated with impaired LTP but not BA hyperexcitability. Interestingly, the middle-aged TgAD rats showed intact short-term extinction and BA LTP, suggestive of a compensatory mechanism, whereas differential neural recruitment in older-aged WT rats may have facilitated short-term extinction. As such, associations between fear extinction memory and amygdala deficits in non-pathological aging and AD are dissociable.

**Significance:** Adults with fear-based disorders like post-traumatic stress disorder are at an increased risk for developing age-related cognitive decline and Alzheimer’s disease (AD). Moreover, negative emotional affect is an early marker of AD. The link between fear-based disorders and AD creates a disadvantage for achieving positive outcomes later in life. Central to the circuitry underlying fear disorders are medial temporal lobe structures like the basal amygdaloid nucleus (BA). However, the role of the BA in fear-based disorders exacerbated by aging and AD is not well understood. Using the TgF344AD rat model of AD, we investigated how fear extinction memory impairments and BA synaptic function are impacted by aging and AD and whether these processes differentially associate in non-pathological aging and AD.

## Introduction

Non-pathological aging is accompanied by impaired executive functions (Buckner, 2004) and a greater prevalence of fear-based neuropsychiatric disorders such as post-traumatic stress disorder (PTSD), general anxiety disorder, and panic disorder (Beaudreau and O’Hara, 2008). Due to the persistent and hyperactive properties of fear memories, symptoms of fear-based disorders can worsen with age (Beaudreau and O’Hara, 2008). Unfortunately, younger individuals who managed to recover from PTSD often have a recurrence in old age (Floyd et al., 2002). Critically, PTSD significantly steepens age-related cognitive decline (Yehuda et al., 2005; Green et al., 2016), associates with greater tau accumulation (Mohamed et al., 2019), and doubles the risk of developing Alzheimer’s disease (AD) and other dementia in older individuals (Qureshi et al., 2010; Yaffe et al., 2010).

Though non-pathological and pathological aging are cognitively and biologically dissociable, aging is the greatest risk factor for the development of neurodegenerative diseases such as AD (Swerdlow, 2007; National Institute on Aging, 2019). Indeed, cognitive and emotional impairments worsened by neuropsychiatric disorders like PTSD in old age may be indicative of an irreversible trajectory toward developing AD. Consistent with this are the findings that individuals with AD have a greater prevalence of fear-based disorders (Burke et al., 2018). Moreover, aversive emotional memories, such as those associated with fear-based neuropsychiatric disorders may be an early marker of AD and dementia (Neary et al., 1998; McKhann et al., 2001; Hoefer et al., 2008), even prior to other measurable cognitive deficits

While the focus of many cognitive aging and AD studies has been on the prefrontal cortex (PFC) and hippocampus (HPC), a unifying thread linking PTSD, cognitive aging, and AD is aberrant activity in the amygdala (Shin et al., 2006; Wright et al., 2007). Topographical studies have established the basal amygdaloid nucleus (BA) of the basolateral amygdala (BLA) complex receives abundant inputs from the PFC and HPC (Mcdonald et al., 1996; McDonald and Mott, 2017). The BLA provides emotional valence (positive or negative) to memory (Garavan et al., 2001; Belova et al., 2008; Beyeler et al., 2016) and serves as an integrative hub in the underlying processes associated with fear memory encoding, consolidation, and retrieval (Maren, 2001; Kochli et al., 2015). Specifically, synaptic communication between the BLA and PFC supports the extinction of fear memories (Peters et al., 2009), whereas communication between BLA and HPC supports fear acquisition and the modulation of fear extinction (Maren and Quirk, 2004; Sierra-Mercado et al., 2011). Critically, the inability to extinguish hyperactive fear memories is a core component of PTSD driven by a hyperexcitable BLA that may be exacerbated by aging and AD (Shin et al., 2006; Wright et al., 2007).

The transgenic Fisher 344 AD (TgAD) rat has been well-characterized as a comprehensive model of AD that develops progressive amyloid pathology, endogenous hyperphosporylated tau, neuroinflammation, and cognitive decline across the life-span (Cohen et al., 2013). Given the recent link between fear-based disorders, cognitive aging, and Alzheimer’s disease (Burke et al., 2018), the current study leveraged this rat model of AD to collectively assess the effects of non-pathological aging and AD on fear extinction memory and BA synaptic function in addition to the relationship between these factors. Indeed, defining the synaptic mechanisms linking these factors is foundational for the development of interventional strategies to remediate poor outcomes in aging and neurodegeneration. As a major step toward this goal, young adult, middle-aged, and older-aged WT and TgAD rats underwent fear conditioning and extinction behavior, followed by extracellular field recording in slices containing the BA. We then employed a principal component analysis (PCA) to determine if associations between fear extinction and BLA function differed by age and AD. In general, our results demonstrate aging in the presence or absence of AD result in unique fear extinction impairments and BA synaptic deficits.

## Materials and Methods

### Subjects

A total of 101 (see Table 1 for sample sizes) wild type (WT) and TgF344-AD (TgAD) rats were used for behavioral and *ex vivo* brain slice electrophysiological experiments. As previously described (Smith and McMahon, 2018; Goodman et al., 2021), transgenic rats (TgAD) harboring the human Swedish mutation amyloid precursor protein (APP^swe^) and the presenilin-1 exon 9 deletion mutant (PS1^ΔE9^) were bred with WT F344 females (Envigo: previously Harlan Laboratories) at the University of Alabama at Birmingham. All breeding and experimental procedures were approved by the University of Alabama at Birmingham Institutional Animal Care and Use Committee and follow guidelines set by the National Institutes of Health. The original breeding pair was obtained from University of Southern California, Los Angeles, CA (Cohen et al., 2013). Of the 101 rats, nineteen 24-mo-old WT were obtained from the National Institute on Aging aged rodent colony maintained by Charles River Laboratories. Rats were maintained under standard animal care facility conditions with food (catalog #Harlan 2916, Teklad Diets) and water *ad libitum* and a 12 h reverse light/dark cycle (lights on at 7am) at 22°C and 50% humidity. Rats were housed in standard rat cages (height, 7 inches; floor area, 144 square inches) in same-sex groups of 4 or less at weights of ∼300 g or two per cage when ≥400 g. Rats were aged from birth to experimental age groups categorized as young adults (YA: 5.67–7.59 mo, average of 6.39 mo), middle-aged (MA: 12.17–14.16 mo, average of 13.36 mo), and older-aged (OA: 22.65–25.13, average of 24.53 mo). While all rats were included in behavioral experiments, only a subset of these rats were assigned to electrophysiological experiments. Prior to contextual fear conditioning, all rats were single housed. Final group sizes for behavioral and electrophysiological experiments are described in Table 1.

**Table 1.** All group sample sizes for behavioral and electrophysiological experiments. A subset of all the rats that received contextual fear conditioning were assigned to electrophysiological experiments, and as such both behavioral and electrophysiological measures were available for each rat.

| Age | Young Adult (YA) |  |  |  | Middle-Aged (MA) |  |  |  | Older-Aged (OA) |  |  |  |
| --- | --- | --- | --- | --- | --- | --- | --- | --- | --- | --- | --- | --- |
| Genotype | WT |  | TgAD |  | WT |  | TgAD |  | WT |  | TgAD |  |
| Sex | M | F | M | F | M | F | M | F | M | F | M | F |
| Behavior (ns) | 8 | 9 | 8 | 7 | 8 | 12 | 11 | 11 | 5 | 16 | 4 | 2 |
| Electrophysiology (ns) | 5 | 4 | 4 | 4 | 4 | 6 | 6 | 5 | 3 | 6 | 3 | 2 |

### Contextual fear conditioning

#### Fear memory acquisition

Day 1 consisted of fear memory acquisition in context A (Figure 1A) which consisted of a custom operant conditioning chamber (29.53 × 23.5 × 20.96 cm, Med Associates) comprised of featureless walls, a metal grated floor that delivered the shock, and an EtOH scent. Prior to testing, all rats were habituated in context-specific holding rooms for a minimum of 15 min, and the transportation route between vivarium and holding room was specific to context A. After a 2min baseline to determine basal freezing behavior, rats received four trials consisting of a 20 s tone (2000Hz at 85Db), serving as the conditioned stimulus (CS), paired with a foot-shock (1mA), serving as the unconditioned stimulus (US), during the last 2 s of the tone (CS and US co-terminated). Each trial was separated by a 20s intertrial interval. After the final CS-US pairing, a 2min post-paring epoch was used to determine fear memory acquisition in what became the “unsafe” context (see Figure 1B for a detailed schematic). As freezing is a response to threat in rodents, fear expression (and by proxy, fear memory) is operationalized as freezing behavior of at least 1 s except for breathing. Freezing activity video was recorded during each session (Video Freeze, Med Associates). Time spent freezing per each epoch (baseline, CS, US, and post-pairings) was divided by the total time of each epoch generate a percentage of time freezing per epoch.

**Figure 1.**
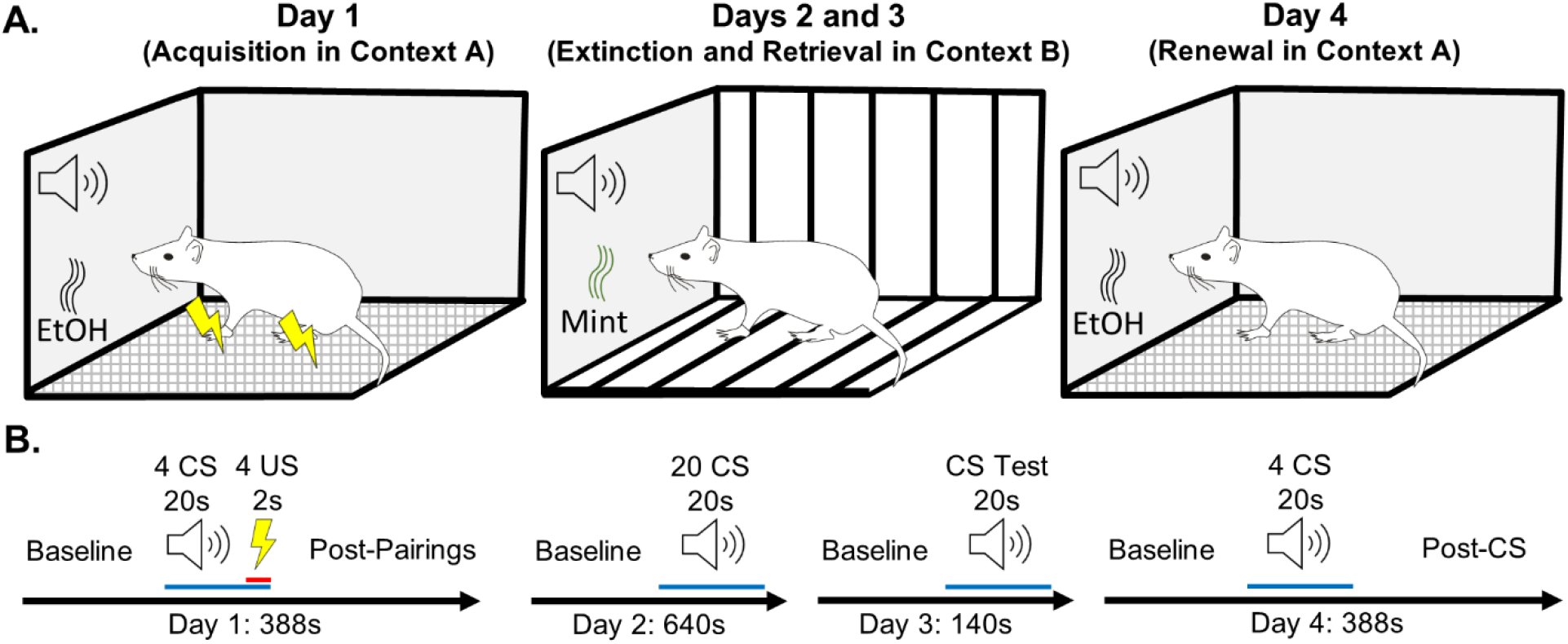
Contextual Fear Conditioning Paradigm. A) Schematic of day-dependent contextual design. B) Trial schematic. Rats progressed through a four-day fear conditioning paradigm in which acquisition in context A took place on day 1, fear extinction in context B took place on day 2, extinction retrieval in context B took place on day 3, and fear renewal in context A took place on day 4. See text in methods for more detail.

**Figure 2.**
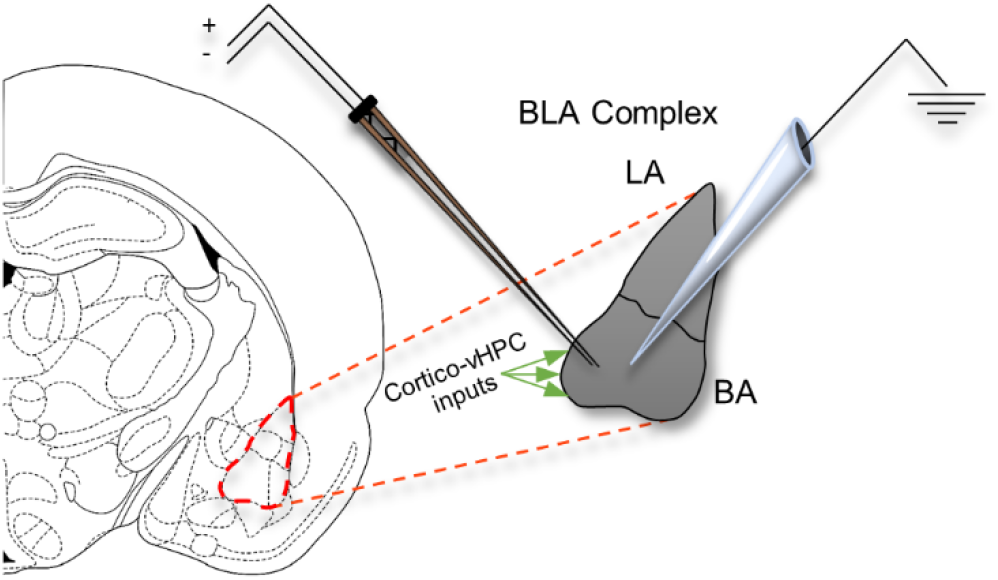
Extracellular recordings targeting the basal amygdaloid nucleus of the BLA complex. Topographical studies of the basolateral amygdala describe abundant cortico-hippocampal inputs to the anterior part, known as the basolateral amygdaloid nucleus, anterior part (BA). To maximize stimulation of these inputs, the stimulating electrode was consistently placed towards the medio-ventral region of the basolateral amygdaloid nucleus anterior part, and the recording electrode was consistently placed between 5 and 10mm lateral relative to the stimulating electrode. LA: Lateral Amygdaloid nucleus of the BLA complex; vHPC: ventral hippocampus.

#### Fear memory extinction

Day 2 consisted of fear memory extinction in context B (Figure 1A). Context B consisted of a different operant conditioning chamber from that used in context A comprised of striped walls, striped plastic flooring that covered the metal grating, and a peppermint scent. Additionally, the holding room during the 15 min habituation period was context-specific, and the transportation route between vivarium and holding room was specific to context B. After a 2 min baseline, rats received 20 trials consisting of the CS (tone) only (i.e., in the absence of the US). Each trial was separated by a 5 s intertrial interval (Figure 1B). As there was no US, context B became the “safe” context.

#### Fear memory extinction retrieval

Day 3 consisted of fear extinction retrieval test (i.e., recall the CS is no longer threatening) in context B (Figure 1A). After a 2 min baseline, rats received 10 trials consisting of the CS only (Figure 1B). Each trial was separated by a 5 s intertrial interval.

#### Fear memory renewal

Day 4 consisted of fear memory renewal, in which rats were placed back into context A (the “unsafe” context) to test if their fear response was renewed (Figure 1A). After a 2 min baseline, rats received four trials consisting of the CS only. Each trial was separated by a 20 s intertrial interval. After the final CS, there was a 2 min post-CS epoch used to determine fear memory renewal to context A (Figure 1B).

### Extracellular field excitatory postsynaptic potential recordings in the BLA

#### BLA slice preparation

Brain slices containing BLA were prepared from rats a minimum of two weeks after contextual fear conditioning. Rats were anesthetized via deep isoflurane inhalation and rapidly decapitated, and brains were rapidly extracted. Coronal slices (400 µm) containing BLA were prepared using a vibratome (model VT1000P, Leica). As per Goodman et al., 2021, slices were made in low Na^+^, sucrose-substituted ice-cold artificial CSF (aCSF) containing (in mM): NaCl (85), KCl (2.5), MgSO_4_ (4), CaCl_2_ (0.5), NaH_2_PO_4_ (1.25), NaHCO_3_ (25), glucose (25), and sucrose (75), saturated with 95% O_2_, 5% CO_2_, at pH 7.4. Slices were held in a water bath at 26°C for 30 min in standard aCSF containing the following (in mM): NaCl (119.0), KCl (2.5), MgSO4 (1.3), CaCl_2_ (2.5), NaH_2_PO_4_ (1.0), NaHCO3 (26), and glucose (11), saturated with 95% O_2_, 5% CO_2_, at pH7.4 before transfer to the submersion chamber for recordings.

#### Stimulus-response (Input/Output I/O)

To determine the strength of basal synaptic transmission, brain slices containing BLA were prepared from rats a minimum of two weeks after contextual fear conditioning. Extracellular field excitatory postsynaptic potentials (fEPSPs) were recorded from BLA slices in a submersion chamber continuously perfused with warm (27–29°C) aCSF containing the GABA_A_ channel blocker, picrotoxin (PTX, 40 μM) to isolate excitatory transmission. Baseline fEPSPs were generated using a bipolar simulating electrode placed within 1 mm of an aCSF-PTX-filled glass recording electrode and by stimulating the medial portion of the basal amygdaloid nucleus (BA; 0.1 Hz for 200 µs) to activate cortical and hippocampal inputs. The fEPSP peak amplitude was measured at increasing stimulus intensities at 5µA steps from 20 to 100µA. Paired pulse stimulation (50 ms interval) was delivered throughout the experiment to assess facilitation or depression.

#### Long-term potentiation (LTP)

Following I/O experiments, the simulation intensity was adjusted to generate approximately 50% of the maximum peak amplitude response and a 10 min baseline was recorded prior to delivering high-frequency stimulation (HFS) to induce long-term potentiation (HFS: 4 trains of 100Hz, 500ms duration, separated by 20s; (Huang and Kandel, 2007; Humeau et al., 2007). The fEPSP peak amplitude was measured, transformed to percent of baseline (%BL), and plotted over time. The LTP magnitude was measured by comparing peak amplitude at 30 min post-tetanus to baseline.

### Statistical analysis and experimental design

#### General statistical approach

Unless otherwise noted, all statistical analyses were performed in SPPSS28 v280.0.0(190). In all analyses, the alpha (α) was set to 0.05, and when Mauchly’s test of sphericity was violated, the Huynh-Feldt p-value correction was applied. When there were significant effects, the effect sizes were reported as η_p_^2^ for ANOVAs and Cohen’s *d* for t-tests. Additionally, observed power for significant effects was reported as 1-β. For brevity, all null effects were reported as consolidated F-statistics (or t-statistics), p-values, and only the lowest and highest values for each were given. Outliers were identified as rats that expressed pre-conditioned freezing during the acquisition baseline epoch using the outlier identification analysis in SPSS. In all analyses, genotype and sex were coded as nominal variables, age was coded as an ordinal variable, and any repeated measure was coded as continuous. All figures were generated in GraphPad Prism v9.3.0(463).

#### Contextual fear conditioning

In general, a hierarchical approach was used in all data analyses. Percent time freezing was initially analyzed using a mixed-factor ANOVA (age × genotype × sex × trial) with age (3 levels: YA, MA, OA), genotype (2 levels: WT and TgAD), and sex (2 levels: male and female) as between-subjects factors, and trial (6 levels for acquisition: BL, CSUS1, CSUS2, CSUS3, CSUS4, and PP; 4 levels for extinction, retrieval, and renewal: ExF, ExL, Ret, Ren; and 3 levels for context: A_1_, B, and A_2_) as the within-subjects factors. While effects of sex are reported, sex is underpowered in older-aged TgAD rats. Therefore, only main effects and interactions between age and genotype were followed up with pairwise comparisons using Fishers Least Significant Difference (LSD). Significant pairwise comparisons were reported as mean difference followed by p-values. A second tier of analysis was used that focused on effects of genotype within an age group by using a two-factor ANOVA (genotype × trial). Finally, to determine if individual groups were successful at acquisition, extinction, retrieval, and renewal, paired-samples t-tests were used for each group separately. For acquisition, paired-samples t-tests compared BL to PP. For extinction and retrieval, paired-samples t-tests compared both the last extinction trial (ExL) and the first retrieval (Ret) trial to the first extinction trial (ExF). For renewal, paired-samples t-tests compared renewal (Ren) to Ret. To compare %time freezing during context testing, paired-samples t-tests were used between context A during acquisition (A_1_) or context A (A_2_) during renewal to context B (after extinction training).

#### Behavioral control measures

To account for the possibility that differences in %time freezing during were confounded by shock perception, motion data was extracted from US epochs, transformed to reflect the relative change in motion across trials (BL to US4), and analyzed with a mixed-factor ANOVA (age × genotype × sex × trial). To further account for the possibility that habituation may confound differences in %time freezing, the relative change in motion between baselines on all days was analyzed using a mixed-factor ANOVA (age × genotype × sex × day).

Finally, the prolonged stress of fear conditioning has been shown to initiate weight loss due to the inhibition of consummatory behavior in rats (Pare, 1965). As an additional measure to account for habituation, each rat’s weight (in grams) was analyzed across days using a multi-factor ANOVA (age × genotype × sex × day).

#### Extracellular fEPSPs

In all electrophysiology experiments, sex was underpowered and therefore excluded as a factor. In the I/O experiments, fEPSP peak amplitude across stimulation intensity was analyzed using a mixed-factor ANOVA (age × genotype × intensity) with age (3 levels) and genotype (2 levels) as between-subjects factors, and intensity as the within-subjects factor (17 levels: 20–100µA in 5µA steps). Main effects, interactions, and a second tier of analyses (genotype × intensity at each age) were followed up as stated above. For each fEPSP recorded in the I/O experiments, a coastline burst index (CBI) was generated to assess possible hyperexcitability (Korn et al., 1987; Stewart et al., 2017; Widman and McMahon, 2018). To leverage the full design of each ANOVA, missing data points for I/O experiments in n=1 young adult WT and n=1 young adult TgAD were computed by linear interpolation using the missing data tool in SPSS (note that effects and interpretations were not influenced by missing data). For LTP experiments, peak amplitude was transformed to percent of baseline, and the initial analysis used a mixed-factor ANOVA (age × genotype × time) with age and genotype as described above and time as the within-subjects factor (40 levels: minutes -1–30). Pairwise comparisons were used as stated above, and a follow-up genotype × time ANOVA was used within each age group as stated above. In addition, to determine if individual groups were successful at short-term potentiation (STP) or LTP, paired-samples t-tests were used to compare peak amplitudes (as %BL) during the first 5 minutes post-HFS to BL (STP) and to compare peak amplitudes (as %BL) at 30 minutes post-HFS to BL (LTP) for each group separately.

#### Extracellular fEPSP control measures

To account for day-to-day variation during I/O experiments, CBIs during the pre-stimulus period from each recording (first 20ms) were compared using a mixed-factor ANOVA (age × genotype × intensity). To account for potential baseline differences confounding effects of age or genotype in LTP experiments, baselines were compared using a mixed-factor ANOVA (age × genotype × time).

#### Principal Components Analysis

For all rats with behavioral and electrophysiological data, a principal components analysis (PCA) was used to analyze associations between fear memory extinction and BA synaptic function. Standardized scores for percent time freezing during MPT, Ext, and Ret were loaded to represent CS memory, extinction, and extinction memory retrieval, respectively. Standardized scores of I/O peak amplitudes and peak amplitudes 30 min post-HFS were loaded to represent synaptic physiology. To avoid split loadings, the rotation used was Varimax with Kaiser normalization. An Eigenvalue >1 was considered meaningful, and factor loadings >0.6 were considered significant. Each component score was extracted as regression loadings, and the unique loadings for each animal were used to plot its distribution upon each component. To determine group differences, regression loadings were analyzed using a multivariate ANOVA (age × genotype) with each component as a dependent variable (DiStefano et al., 2009). Finally, using a combination of SPSS and R (ggplot2), regression loadings were utilized to visualize group clustering within the rotated space, and 95% confidence ellipses were generated utilizing the ellipse stat function.

## Results

### Effects of age and AD on fear memory

#### Fear memory acquisition

We first tested whether differences exist between WT and TgAD rats in their ability to acquire fear memory over the lifespan. On day 1, young adult, middle-aged, and older-aged WT and TgAD rats received CS-US pairings after a baseline period in context A. Two outliers (n=1 young adult WT rat; n=1 middle-aged TgAD rat) were identified and removed from this and all subsequent analyses. Aging affected %time freezing (F_(2, 87)_=9.765, p<0.001, η_p_^2^=0.183, 1-β=0.980) such that middle-aged rats froze significantly less compared to young (−11.460, p<0.001) and older rats (−9.063; p=0.008); all TgAD rats froze less (−9.447) relative to WT (F_(1, 87)_=12.979, p<0.001, η_p_^2^=0.130, 1-β=0.945); all males froze less (−6.005) relative to females (F_(1, 87)_=5.243, p=0.024, η_p_^2^=0.057, 1-β=0.620); and all rats froze more in all trials after CS-US1 relative to baseline (+14.676–25.763; F_(5, 435)_=22.638, p<0.001, η_p_^2^=0.206, 1-β=1.000). Furthermore, there were significant age × genotype (F_(2, 87)_=5.093, p=0.008, η_p_^2^=0.105, 1-β=0.809), age × trial (F_(10, 435)_=2.369, p=0.013, η_p_^2^=0.052, 1-β=0.918), and genotype × trial (F_(5, 435)_=2.535, p=0.033, η_p_^2^=0.028, 1-β=0.756) interactions. All other interactions were non-significant (Fs_(1–10, 87–435)_=0.210–2.746, ps=0.101–0.957). See Figure 3A.

**Figure 3.**
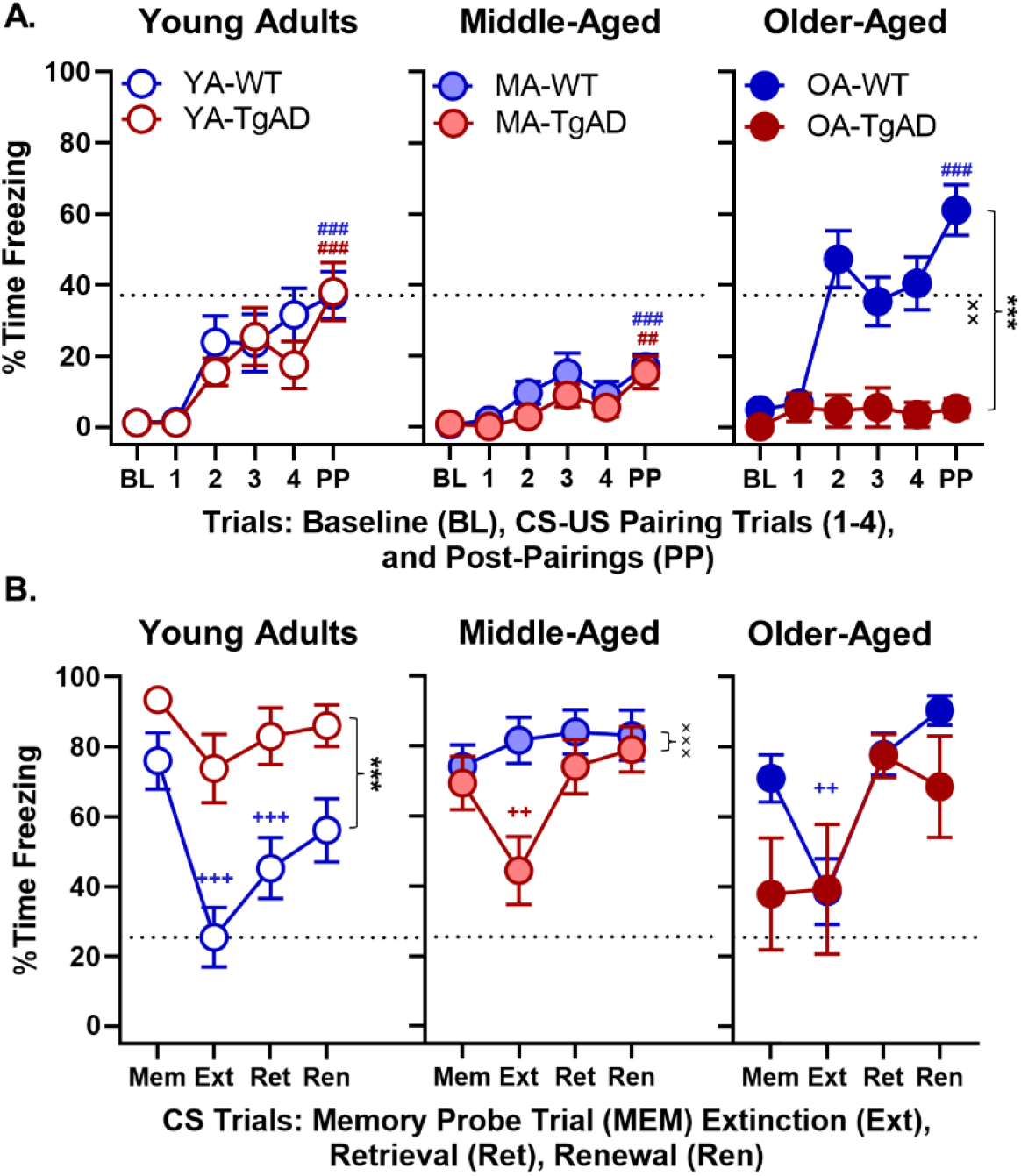
Acquisition, extinction, retrieval, and renewal. **A)** Day 1 acquisition. There were no genotype differences within the young adult or middle-aged groups. Within the older-aged group, there was a significant main effect of genotype and significant genotype by trial interaction such that WT showed greater %time freezing relative to TgAD. Within the exception of older-aged TgAD rats, all groups acquired fear memory during day 1. The x-axis represents baseline (BL), CS-US paring trials (1-4), and post-pairings (PP). **B)** Days 2-4 extinction, retrieval, and renewal. Within the young adults, there was a main effect of genotype such that TgAD rats showed greater %time freezing relative to WT. While young adult WT rats showed successful fear extinction and retrieval, young adult TgAD rats did not. Within the middle-aged group, there was a significant genotype by trial interaction. Middle-aged TgAD rats showed intact acute fear extinction but not retrieval, whereas the WT middle-aged rats did not show either. There were no genotypic differences within the older-aged group. Older-aged WT rats showed intact acute fear extinction but not retrieval, whereas there was no extinction or retrieval in the older-aged TgAD rats. Data are plotted as the means with SEM. ***p<0.001 for main effects of genotype; ^××^p<0.01, ^×××^p<0.001 for genotype × intensity interactions; ^++^p<0.01, ^+++^p<0.001 for significant differences between Ext or retrieval and MPT within groups (blue for WT and red for TgAD); ^##^p<0.01, ^###^p<0.001 for significant differences between BL and PP during acquisition within groups (blue for WT and red for TgAD). Dotted black line represents mean of YA-WT PP in A and mean of YA-WT Ext in B.

To better define the nature of these interactions and to test if the degree of acquisition was different between WT and TgAD rats at each age, we followed up with a genotype × trial ANOVA. Within the young adult age group, genotypes were not different in any trial (Fs_(1–5, 29–145)_=0.407– 0.813, ps=0.509–0.528). However, all young adults progressively froze more after baseline (trial: F_(5, 145)_=16.352, p<0.001, η_p_^2^=0.361, 1-β=1.000; Figure 3A). Similarly, within the middle-aged group, genotypes were not different at any trial (Fs_(1–5, 39–195)_=0.546–1.556, ps=0.220–0.673), and all rats progressively froze more after baseline (trial: F_(5, 195)_=10.727, p<0.001, η_p_^2^=0.216, 1-β=1.000; Figure 3A). In contrast, older-aged TgAD rats froze less after baseline relative to their WT counterparts (genotype: F_(1, 25)_=21.507, p<0.001, η_p_^2^=0.462, 1-β=0.994; genotype × trial: F_(5, 125)_=3.910, p=0.003, η_p_^2^=0.135, 1-β=0.928; trial: F_(5, 125)_=4.668, p<0.001, η_p_^2^=0.157, 1-β=0.966; Figure 3A).

Finally, to confirm that all WT and TgAD rats acquired fear memory, we used paired-samples t-tests to compare %time freezing during baseline to %time freezing during post-pairings. Figure 3A shows increased freezing from baseline to post-pairings, confirming fear memory acquisition in young adult WT rats (t_(15)_=5.476, p<0.001, *d*=1.369), young adult TgAD rats (t_(14)_=4.492, p<0.001, *d*=1.160), middle-aged WT rats (t_(19)_=4.611, p<0.001, *d*=1.031), middle-aged TgAD rats (t_(20)_=3.082, p=0.006, *d*=0.672), older-aged WT rats (t_(20)_=8.168, p<0.001, *d*=1.782), but not older-aged TgAD rats (t_(5)_=1.932, p=0.111, *d*=0.789).

Collectively, these results suggest varying degrees of intact fear memory acquisition in all rats except older-aged TgAD rats. It should be notated that, in contrast to previous reports (Maren et al., 1994; Pryce et al., 1999), we found freezing time during acquisition was modestly greater in females, which is consistent with females being more sensitive to punishment (Miettunen et al., 2007; Orsini et al., 2016; Liley et al., 2019; Hernandez et al., 2020). Furthermore, female 3×TgAD and control mice have greater freezing relative to males (Stover et al., 2015). Notably, these sex differences do not explain the age and genotype effects on acquisition.

#### Fear memory extinction, retrieval, and renewal

We then tested whether fear memory of the CS was intact within-groups on day 2 by using paired-samples t-tests to compare %time freezing between baseline (Table 2) and the CS memory probe trial (Mem). These results revealed young adult WT (t_(15)_=7.779, p<0.001, *d*=1.945), young adult TgAD (t_(14)_=4.993, p<0.001, *d*=1.289), middle-aged WT (t_(19)_=5.808, p<0.001, *d*=1.299), middle-aged TgAD (t_(20)_=6.687, p<0.001, *d*=1.459), and older-aged WT rats (t_(20)_=7.669, p<0.001, *d*=1.673) increased freezing suggesting robust expression of fear memory to the CS. The older-aged TgAD rats did not show an increase in %time freezing between baseline and the probe CS memory trial (t_(5)_=0.679, p=0.527). However, to address whether the null effect was confounded by an increase in %time freezing during baseline, a comparison of %time freezing between the final CS trial during acquisition and MPT revealed a significant increase in fear memory expression (t_(5)_=2.044, p=0.048, *d*=0.834).

**Table 2.**
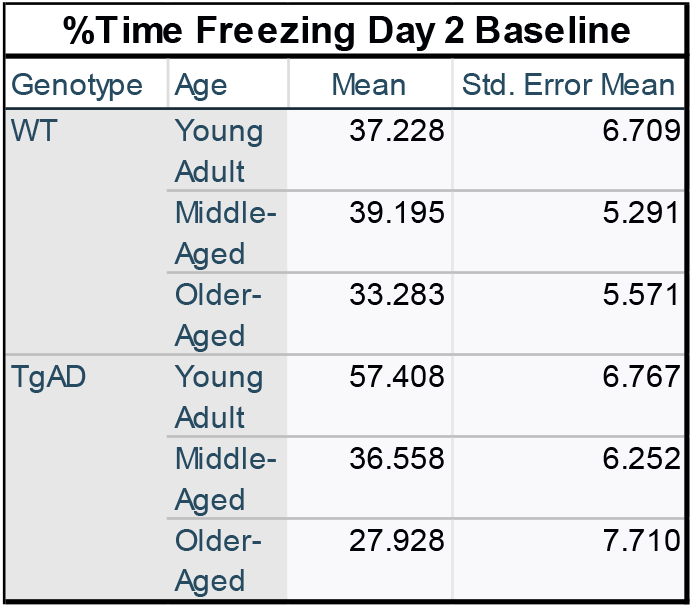
Percent time freezing during baseline on day 2.

After confirming there was fear memory in all groups, we tested whether differences exist between WT and TgAD rats over the lifespan in fear memory extinction, retrieval, and renewal. Rats received CS trials without the US pairing in a novel context B to induce extinction on day 2. Then, we tested fear extinction memory retrieval in WT and TgAD rats over the lifespan on day 3. Finally, all rats were given CS trials without US pairings in context A to test for fear memory renewal on day 4. While an analysis of %time freezing during extinction, extinction memory retrieval, and renewal using a mixed-factor ANOVA (age × genotype × sex × trial) did not reveal any main effects of age, genotype, or sex (Fs_(1–2, 87)_=0.000–1.549, ps=0.218–0.996), the main effect of trial was significant (F_(3, 261)_=21.423, p<0.001, η_p_^2^=0.198, 1-β=1.000). Furthermore, there were significant age × genotype (F_(2, 87)_=9.529, p<0.001, η_p_^2^=0.180, 1-β=0.977), age × trial (F_(6, 261)_=5.724, p<0.001, η_p_^2^=0.116, 1-β=0.997), and age × genotype × trial (F_(6, 261)_=4.219, p<0.001, η_p_^2^=0.088, 1-β=0.978) interactions (Figure 3B). All other interactions were non-significant (Fs_(1–6, 87–261)_=0.382–2.128, ps=0.097–0.766).

To better understand the interaction, we tested if freezing during extinction, retrieval, and renewal differed between WT and TgAD groups at each age. Within the young adult age group, TgAD rats froze more overall relative to WT (genotype: F_(1, 29)_=16.523, p<0.001, η_p_^2^=0.363, 1-β=0.975; genotype × trial: F_(3, 87)_=2.128, p=0.103; trial: F_(3, 87)_=10.723, p<0.001, η_p_^2^=0.270, 1-β=0.999; Figure 3B). Within the middle-aged group, genotype interacted with trial (F_(3, 117)_=6.106, p<0.001, η_p_^2^=0.135, 1-β=0.956) such that WT rats froze more (+37.229, p=0.003) only during Ext relative to TgAD (trial: F_(3, 117)_=6.638, p<0.001, η_p_^2^=0.145, 1-β=0.970; genotype: F_(1, 39)_=2.534, p=0.120; Figure 3B). In the older-aged group, genotypes were not different at any trial (Fs_(1–3, 25– 75)_=1.632–1.915, ps=0.179–0.189), and all rats froze more during renewal testing (F_(3, 75)_=9.013, p<0.001, η_p_^2^=0.265, 1-β=0.994).

We then tested if WT and TgAD rats expressed fear memory extinction, retrieval, and renewal across the lifespan with paired-samples t-tests. Young adult WT rats decreased freezing during extinction (t_(15)_=-5.294, p<0.001, *d*=1.324; Figure 3B) and retrieval relative to the memory probe trial (t_(15)_=-4.529, p<0.001, *d*=1.132) but not during renewal relative to retrieval (t_(15)_=1.427, p=0.174), suggesting there was intact extinction and extinction memory retrieval, whereas the lack of renewal may indicate some resilience to expressing fear that as a result of maintained fear extinction memory. Young adult TgAD rats only tended to slightly decrease freezing during extinction (t_(14)_=-2.095, p=0.055, *d*=0.541; Figure 3B) with no other differences (ts_(14)_= 0.488– 1.110, ps=0.286–0.633), suggesting an early life impairment in fear extinction. Middle-aged WT rats showed no differences between trials (ts_(19)_= 0.170–1.590, ps=0.128–0.867). Surprisingly, middle-aged TgAD rats did decrease freezing during extinction (t_(20)_=-2.881, p=0.009, *d*=0.629; Figure 3B) but not retrieval and showed no other differences (ts_(20)_= 0.873–1.351, ps=0.192– 0.393), suggesting a compensation in the ability to extinguish fear memory but not the ability to maintain that memory long-term. Older-aged WT rats also decreased freezing during extinction (t_(20)_=-3.077, p=0.006, *d*=0.671; Figure 3B) but not retrieval and showed no other differences (ts_(20)_= 1.324–1.741, ps=0.097–0.200), suggesting a late-life reemergence in the ability to extinguish fear memory as a function of non-pathological aging without maintenance of that extinction memory. Finally, older-aged TgAD rats showed no differences between trials (ts_(5)_= 0.066–2.420, ps=0.060–0.950).

These results show successful fear memory extinction and retrieval in the absence of fear renewal in the young adult WT rats, whereas the young adult TgAD rats were extinction impaired. Moreover, while middle-aged WT rats were extinction impaired, middle-aged TgAD rats surprisingly showed acute fear memory extinction, potentially resulting from an age-dependent compensatory mechanism that is not sustained into old age. However, middle-aged TgAD rats were impaired in the ability to retrieve fear extinction memory after a 24-hr delay. Similarly, while the older-aged WT rats showed acute fear memory extinction, they were impaired at retrieval. Fear memory expression in older-aged TgAD rats did emerge during the memory probe trial, but instead of extinguishing fear memory, they showed increased fear memory expression during retrieval and renewal testing. The absence of fear renewal may be due to a ceiling effect in fear expression during retrieval. In the young adult WT, however, no significant renewal to the CS may be explained by either resilience against fear renewal or by slightly elevated fear expression during retrieval. As there was a numerical increase in fear memory expression from retrieval to renewal, it is more likely the null effect is due to original fear memory savings during retrieval, and given more extinction training, it is possible the difference in fear memory expression between retrieval and renewal would become significant in young WT rats.

#### Fear memory to context

As fear memory to context and CS are dissociable, in a separate analysis, we tested if freezing to contextual cues alone differed between groups. While statistical analyses did not reveal any main effects of age, genotype, or sex (Fs_(1–2, 87)_=0.090–1.372, ps=0.245–0.860), the main effect of trial was significant (F_(2, 174)_=68.957, p<0.001, η_p_^2^=0.442, 1-β=1.000). Furthermore, there were age × genotype (F_(2, 87)_=7.473, p=0.001, η_p_^2^=0.147, 1-β=0.935, age × trial (F_(4, 174)_=6.854, p<0.001, η_p_^2^=0.136, 1-β=0.993), genotype × trial (F_(2, 174)_=5.437, p=0.005, η_p_^2^=0.059, 1-β=0.842), and genotype × sex × trial (F_(2, 174)_=3.890, p=0.022, η_p_^2^=0.043, 1-β=0.697) interactions. No other interactions were significant (Fs_(1–4, 87–174)_=0.016–1.854, ps=0.132–0.984).

We then tested if freezing to context differed between genotype groups at each age in a follow-up analysis. Within the young adults WT froze less during context B and A_2_ testing relative to their TgAD counterparts (genotype: F_(1, 29)_=6.306, p=0.018, η_p_^2^=0.179, 1-β=0.680; trial: F_(2, 58)_=13.632, p<0.001, η_p_^2^=0.320, 1-β=0.997; genotype × trial: F_(2, 58)_=4.898, p=0.011, η_p_^2^=0.144, 1-β=0.785; Figure 4), consistent with the interpretation that TgAD rats generalized contexts. Within the middle-aged group, there were no genotypic differences at any trial (Fs_(1–2, 39–78)_=0.048–1.101, ps=0.338–0.828), but all rats progressively froze more across context testing (F_(2, 78)_=57.083, p<0.001, η_p_^2^=0.144, 1-β=0.785). Within the older-aged group, WT rats froze more during all of context testing (genotype: F_(1, 25)_=12.930, p=0.001, η_p_^2^=0.341, 1-β=0.932; trial: F_(2, 50)_=21.705, p<0.001, η_p_^2^=0.465, 1-β=1.000; genotype × trial: F_(2, 50)_=5.194, p=0.009, η_p_^2^=0.172, 1-β=0.806; Figure 4).

**Figure 4.**
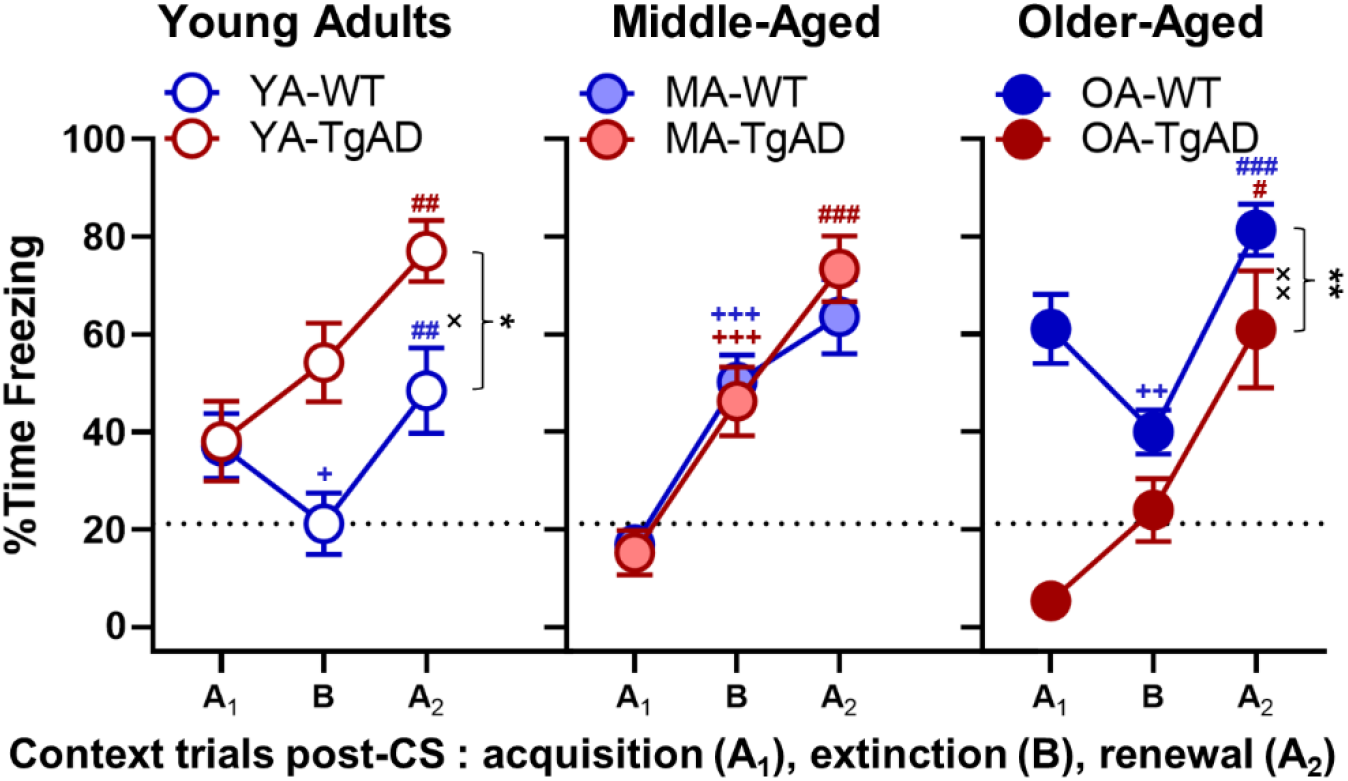
Context. Percent time freezing to different contextual conditions (context A post-pairings during acquisition → context B → context A post-CS during renewal) within each age group. Within the young adults, there was a significant main effect of genotype and a significant genotype by trial interaction such that TgAD rats progressively increased %time freezing relative to WT rats. Within the young adult WT rats, there was decreased %time freezing from context A during acquisition to context B, and then increased freezing from context B to context A during renewal. Within the young adult TgAD rats, there was a significant increase in %time freezing from context B to context A during renewal. There were no genotypic differences in the middle-aged group. Within both WT and TgAD middle-aged rats, there was a progressive increase in %time freezing across context testing, however, the increase between context B and context A during renewal in middle-aged WT rats was not significant. Within the older-aged group, there was a significant main effect of genotype and a significant genotype by trial interaction such that freezing was greater in WT rats relative to TgAD. Older-aged WT rats showed a decrease in %time freezing between context A during acquisition and context B, and an increase between context B and context A during renewal. In contrast, older-aged TgAD rats showed a significant increase in %time freezing between context B and context A during renewal. Data are plotted as the means with SEM. *p<0.05, **p<0.01 for main effects of genotype; ^×^p<0.05, ^××^p<0.01 for genotype × intensity interactions; ^+^p<0.05, ^++^p<0.01, ^+++^p<0.001 for significant differences between context A during acquisition and context B within groups (blue for WT and red for TgAD); ^##^p<0.01, ^###^p<0.001 for significant differences between context B and context A during renewal within groups. Dotted black line represents mean of YA-WT PP for Context B.

Finally, we tested if WT and TgAD rats at all ages could discriminate between “safe” and “unsafe” contexts with paired-samples t-tests. Young adult WT rats froze less to context B relative to context A_1_ (t_(15)_=-2.135, p=0.0496, *d*=-0.534; Figure 4) indicating less fear for context B, and when they were placed back into context A_2_ during renewal, there was an expected freezing increase relative to context B (t_(15)_=3.669, p=0.002, *d*=0.917; Figure 4) demonstrating fear renewal to the “unsafe” context. In contrast, young adult TgAD rats showed a non-significant freezing increase from context A_1_ to B (t_(14)_=1.789, p=0.095) and significant increase from B to A_2_ (t_(15)_=3.194, p=0.007, *d*=0.825; Figure 4) consistent with the inability to discriminate “safe” and “unsafe” contextual cues. Similarly, both middle-aged WT and TgAD rats showed progressive increases in freezing from context A_1_ to context B (WT: t_(19)_=5.925, p<0.001, *d*=1.325; TgAD: t_(20)_=4.090, p<0.001, *d*=0.893; Figure 4) and from context B to context A_2_ (WT: t_(19)_=2.057, p=0.054, *d*=0.460; TgAD: t_(20)_=4.522, p<0.001, *d*=0.987; Figure 4). Older-aged WT rats decreased freezing between context A_1_ and context B (t_(20)_=-2.960, p=0.008, *d*=-0.646; Figure 4) and then increased freezing between context B and context A_2_ (t_(20)_=7.741, p<0.001, *d*=1.630; Figure 4) suggesting the reemergence of some ability to discriminate contextual cues. However, older-aged WT rats expressed greater fear during the “safe” context relative to their young counterparts (t_(35)_=2.493, p=0.018, *d*=0.827) indicating aging is associated with maladaptive responses in emotional regulation. In contrast, older-aged TgAD rats tended to increase freezing between context A_1_ and context B (t_(5)_=2.374, p=0.064, *d*=0.969) and significantly freezing between context B and context A_2_ (t_(5)_=3.660, p=0.015, *d*=1.494; Figure 4). Together, these data suggest context discernment and renewed fear memory is intact in young adult WT rats, whereas all other groups show greater freezing and impairments such that unsafe and safe context were grossly generalized.

#### Behavioral control measures

To determine if shock perception (reactivity) partially explained any effects of age and genotype on freezing across testing phases, we analyzed the change in motion across trials (as defined in the methods). There was an expected increase in motion between baseline and US1 that maintained across successive trials (F_(3, 261)_=132.407, p<0.001, η_p_^2^=0.603, 1-β=1.000), and females responded slightly more (0.299 ±0.145 in pixel change) than males (sex: F_(1, 87)_=4.228, p=0.043, η_p_^2^=0.046, 1-β=0.529; sex × trial: F_(3, 261)_=7.055, p=0.001, η_p_^2^=0.075, 1-β=0.929). However, as noted above, sex as a biological factor did not explain age and genotype effects on extinction, extinction memory retrieval, and renewal. Importantly, there were no effects of age, genotype, or any other interactions on shock perception (Fs_(1–6, 87– 261)_=0.056–2.036, ps=0.157–0.866). As such, differences in shock perception do not explain effects of age and genotype on fear extinction, retrieval, and renewal (Table 3).

**Table 3.** Behavioral control measures. Mean and standard error of each measure for young, middle-aged, and older-aged WT and TgAD rats.

|  |  |  | Genotype |  |  |  |
| --- | --- | --- | --- | --- | --- | --- |
|  |  |  | WT |  | TgAD |  |
| Measure | Age | Independent Variable | Mean | Std. Error | Mean | Std. Error |
| Change in Shock Reactivity | Young Adult | CSUS1 - BL | 3.756 | 0.544 | 3.894 | 0.563 |
|  |  | CSUS2 - CSUS1 | -0.138 | 0.065 | -0.041 | 0.067 |
|  |  | CSUS3 - CSUS2 | 1.127 | 0.432 | -0.037 | 0.447 |
|  |  | CSUS4 - CSUS3 | -0.042 | 0.072 | 0.135 | 0.075 |
|  | Middle-Aged | CSUS1 - BL | 4.582 | 0.496 | 3.305 | 0.475 |
|  |  | CSUS2 - CSUS1 | -0.098 | 0.059 | -0.099 | 0.057 |
|  |  | CSUS3 - CSUS2 | 0.132 | 0.394 | 0.031 | 0.377 |
|  |  | CSUS4 - CSUS3 | 0.022 | 0.066 | 0.073 | 0.063 |
|  | Older-Aged | CSUS1 - BL | 4.101 | 0.557 | 3.759 | 0.942 |
|  |  | CSUS2 - CSUS1 | 0.047 | 0.067 | -0.152 | 0.113 |
|  |  | CSUS3 - CSUS2 | 0.022 | 0.442 | 0.123 | 0.748 |
|  |  | CSUS4 - CSUS3 | -0.094 | 0.074 | -0.064 | 0.125 |
| Change in Baseline Motion | Young Adult | Day2 - Day1 | -0.572 | 0.060 | -0.825 | 0.062 |
|  |  | Day3 - Day2 | 0.751 | 4.850 | 15.013 | 5.020 |
|  |  | Day4 - Day3 | -0.007 | 0.718 | 3.464 | 0.743 |
|  | Middle-Aged | Day2 - Day1 | -0.838 | 0.054 | -0.669 | 0.052 |
|  |  | Day3 - Day2 | -0.170 | 4.428 | -0.484 | 4.238 |
|  |  | Day4 - Day3 | 0.430 | 0.655 | 0.436 | 0.627 |
|  | Older-Aged | Day2 - Day1 | -0.685 | 0.061 | -0.685 | 0.103 |
|  |  | Day3 - Day2 | -0.165 | 4.970 | 0.047 | 8.401 |
|  |  | Day4 - Day3 | 0.125 | 0.735 | -0.123 | 1.243 |
| Body Weights (g) | Young Adult | Day 1 | 356.625 | 8.570 | 368.902 | 8.871 |
|  |  | Day 2 | 354.000 | 8.543 | 367.598 | 8.843 |
|  |  | Day 3 | 354.188 | 8.621 | 365.000 | 8.923 |
|  |  | Day 4 | 352.875 | 8.657 | 364.429 | 8.961 |
|  | Middle-Aged | Day 1 | 422.792 | 7.824 | 432.573 | 7.489 |
|  |  | Day 2 | 420.833 | 7.799 | 431.059 | 7.466 |
|  |  | Day 3 | 419.813 | 7.869 | 431.455 | 7.533 |
|  |  | Day 4 | 418.396 | 7.903 | 431.295 | 7.565 |
|  | Older-Aged | Day 1 | 360.369 | 8.782 | 412.750 | 14.844 |
|  |  | Day 2 | 359.213 | 8.754 | 409.125 | 14.797 |
|  |  | Day 3 | 356.819 | 8.833 | 408.000 | 14.931 |
|  |  | Day 4 | 356.013 | 8.871 | 404.500 | 14.995 |

**Table 4.**
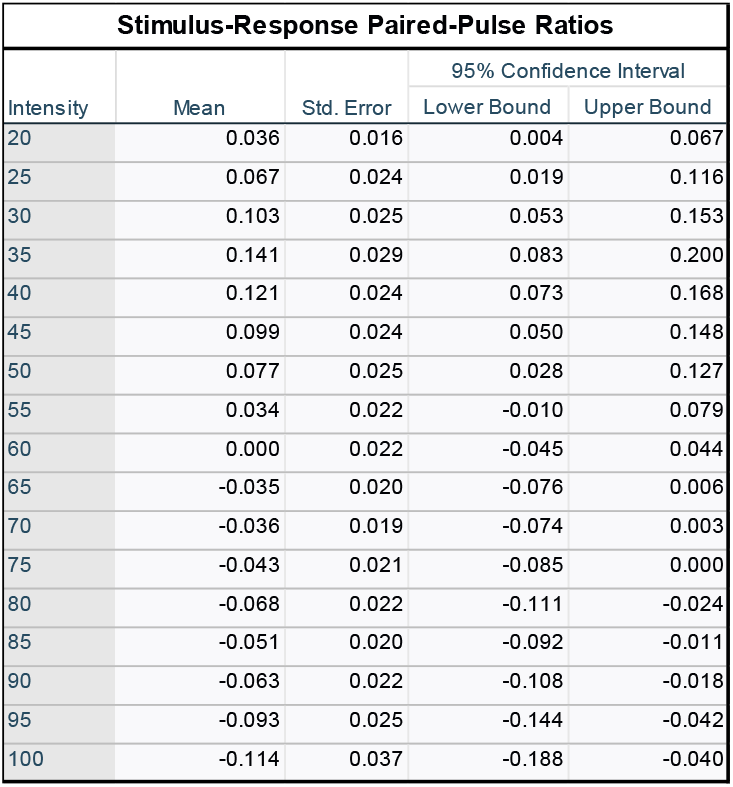
Stimulus-response paired pulse ratios. Mean and standard error of the PPR for each stimulation intensity irrespective of age and genotype.

| Stimulus-Response Paired-Pulse Ratios |  |  |  |  |
| --- | --- | --- | --- | --- |
| Intensity | Mean | Std. Error | 95% Confidence Interval |  |
|  |  |  | Lower Bound | Upper Bound |
| 20 | 0.036 | 0.016 | 0.004 | 0.067 |
| 25 | 0.067 | 0.024 | 0.019 | 0.116 |
| 30 | 0.103 | 0.025 | 0.053 | 0.153 |
| 35 | 0.141 | 0.029 | 0.083 | 0.200 |
| 40 | 0.121 | 0.024 | 0.073 | 0.168 |
| 45 | 0.099 | 0.024 | 0.050 | 0.148 |
| 50 | 0.077 | 0.025 | 0.028 | 0.127 |
| 55 | 0.034 | 0.022 | -0.010 | 0.079 |
| 60 | 0.000 | 0.022 | -0.045 | 0.044 |
| 65 | -0.035 | 0.020 | -0.076 | 0.006 |
| 70 | -0.036 | 0.019 | -0.074 | 0.003 |
| 75 | -0.043 | 0.021 | -0.085 | 0.000 |
| 80 | -0.068 | 0.022 | -0.111 | -0.024 |
| 85 | -0.051 | 0.020 | -0.092 | -0.011 |
| 90 | -0.063 | 0.022 | -0.108 | -0.018 |
| 95 | -0.093 | 0.025 | -0.144 | -0.042 |
| 100 | -0.114 | 0.037 | -0.188 | -0.040 |

We then wanted to determine if habituation partially explained any effects of age and genotype on freezing across testing phases. Therefore we analyzed the relative change in motion in baselines across days and confirmed no main effects or interactions (Fs_(1–4, 87–174)_=0.784–1.670, ps=0.199–0.475). Moreover, an analysis of body weight across days revealed all rats consistently lost weight from day 1 to day 4 (F_(3, 261)_=25.645, p<0.001, η_p_^2^=0.228, 1-β=1.000) suggesting a stress-driven inhibition of consummatory behavior during contextual fear conditioning (Pare, 1965); Table 2). Additionally, TgAD rats weighed more than WT (F_(1, 87)_=9.578, p=0.003, η_p_^2^=0.099, 1-β=0.864; TgAD: 402.22g, ±6.282; WT: 377.66g, ±4.851). An age effect (F_(2, 87)_=33.017, p<0.001, η_p_^2^=0.431, 1-β=1.000) revealed young adults weighed less relative to other age groups (young adults: 360.452g, ±6.167; middle-aged: 426.027g, ±5.415; older-aged: 383.348g, ±8.624; ps=0.034-<0.001). Finally, females expectedly weighed less than males (sex: F_(1, 87)_=456.256, p<0.001, η_p_^2^=0.840, 1-β=1.000; males: 474.706g, ±5.468; females: 305.179g, ±5.757; interactions with sex: Fs_(1–3, 87–261)_=3.143–15.131, ps=0.031–<0.001, η^2^s=0.035-0.184, 1-βs=0.687-0.980). No other interactions were significant (Fs_(2–6, 87–261)_=0.954–2.087, ps=0.072– 0.450). The null effects on change in motion across days coupled with a day-dependent decrease in weight as a marker of stress suggest the effects of age and genotype on %time freezing during fear extinction, retrieval, and renewal were not explained by habituation (Table 3).

### Effects of age and AD on basolateral amygdala synaptic physiology

#### Basal Synaptic Strength

Lesion and *in vivo* electrophysiology studies show the BLA supports the associative learning between CS and US necessary for fear memory acquisition (LeDoux et al., 1990; Romanski et al., 1993). More specifically, localized lesions or protein synthesis blockage in the lateral amygdaloid nucleus (LA) impairs delay and contextual fear memory consolidation, whereas blocking protein synthesis in the BA prevents contextual fear memory consolidation (Kochli et al., 2015). Importantly, BA lesions do not block fear memory acquisition whereas LA lesions do (Amorapanth et al., 2000). As acquisition was not grossly impaired by age or genotype but fear extinction memory was, the behavioral differences observed could be explained by changes in synaptic function at excitatory synapses in the BA nucleus. Thus, we asked whether differences exist in the strength of basal excitatory synaptic transmission in the BA of WT and TgAD rats by recording extracellular fEPSPs in acute slices and performing stimulus response curves (Input/Output (I/O) curves). We measured peak fEPSP amplitude while increasing the stimulus intensity at 5µA increments. While peak fEPSP amplitudes did not differ by age (F_(2, 46)_=1.787, p=0.179), peak fEPSP amplitudes in all TgAD rats were significantly greater relative to all WT (+0.091mV, F_(1, 46)_=11.987, p=0.001, η_p_^2^=0.207, 1-β=0.924; Figure 5B), and all peak amplitudes increased at higher stimulation intensities as expected (F_(16, 736)_=98.820, p<0.001, η_p_^2^=0.682, 1-β=1.000). Although, there were significant age × genotype (F_(2, 46)_=1.787, p=0.049, η_p_^2^=0.123, 1-β=0.585) and genotype × intensity (F_(16, 736)_=3.304, p=0.0496, η_p_^2^=0.067, 1-β=0.564) interactions, no other interactions were significant (Fs_(2–32, 46–736)_=0.711–0.856, ps=0.479–0.565). We followed up this analysis by testing if the I/O curves differed by genotype at each age.

**Figure 5.**
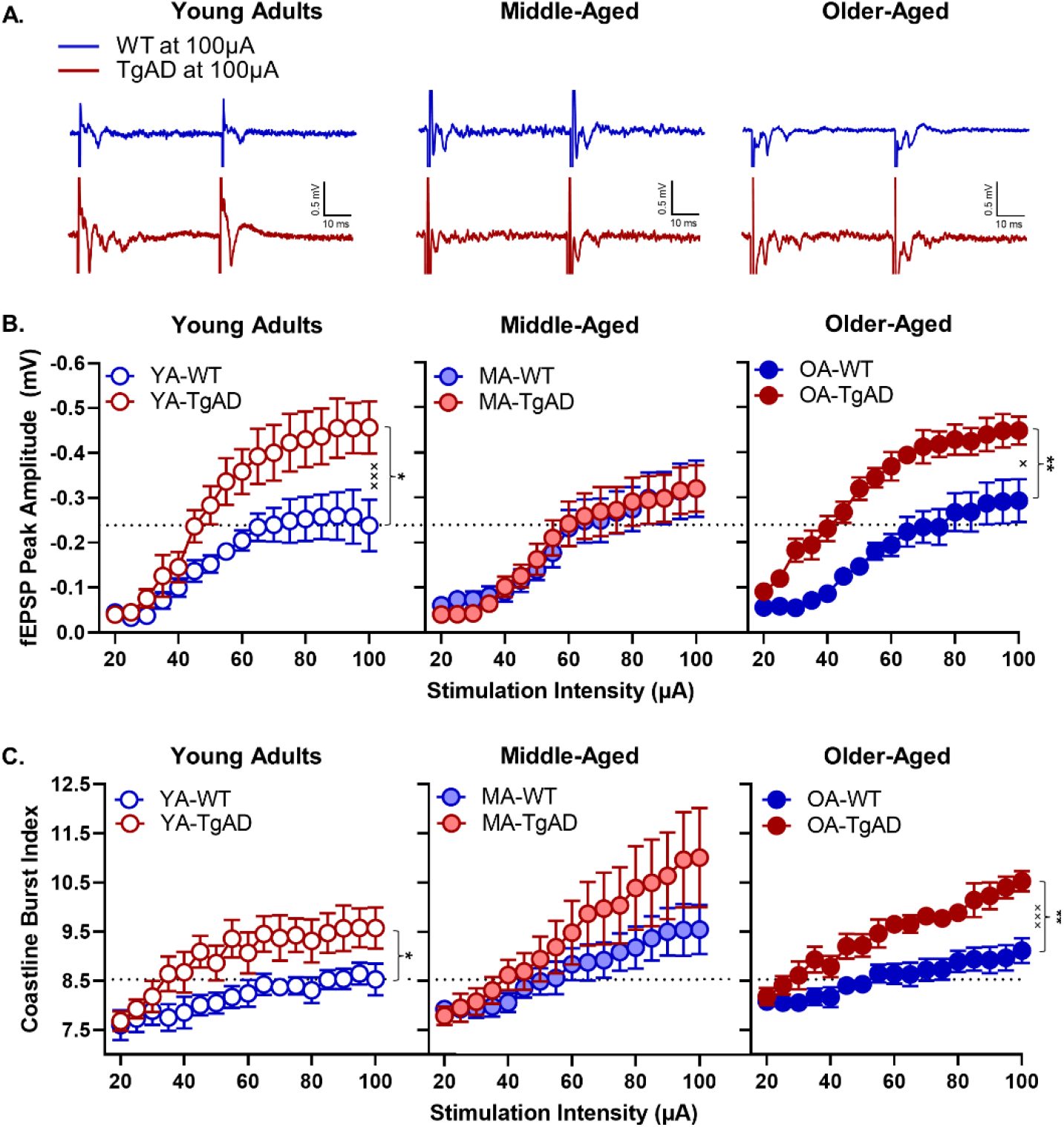
fEPSP peak amplitude responses to increasing stimulation intensity. **A)** Representative traces for each group at maximum simulation intensity (100µA). **B)** fEPSP peak amplitudes within age groups. Within the young adults, there was a significant main effect of genotype and a significant genotype by intensity interaction such that TgAD rats had greater peak amplitudes at higher stimulation intensities relative to WT. There were no differences between WT and TgAD rats within the middle-aged group. Within the older-aged group, there was a significant main effect of genotype and a significant genotype by intensity interaction such that TgAD rats had greater peak amplitudes at higher stimulation intensities relative to WT. There was also a significant age by genotype interaction. **C)** The coastline burst index for each age group. Within the young adults, there was a significant main effect of genotype such that TgAD rats had greater BA activity. There were no genotypic differences among the middle-aged rats. Within the older-aged group, there was a significant main effect of genotype and a significant genotype by intensity interaction such that older-aged TgAD rats had greater activity at higher stimulation intensities relative to WT rats. There was also a trending age by intensity interaction irrespective of genotype. Data are plotted as the means with SEM. *p<0.05, **p<0.01 for main effects of genotype; ×p<0.05, ×××p<0.001 for genotype × intensity interactions. Dotted black line represents mean of YA-WT at 100µA.

Within the young adults, TgAD rats showed larger peak amplitudes at higher intensities relative to WT (genotype: F_(1, 15)_=8.017, p=0.013, η_p_^2^=0.348, 1-β=0.754; intensity: F_(16, 240)_=34.426, p<0.001, η_p_^2^=0.697, 1-β=1.000; genotype × intensity: F_(16, 240)_=2.952, p<0.001, η_p_^2^=0.164, 1-β=0.998; Figure 5B). Within the middle-aged group, there were no genotypic differences (Fs_(1–16, 19–304)_=0.002–0.268, ps=0.708–0.962), but peak amplitudes increased at higher stimulation intensities in all rats (F_(16, 304)_=32.046, p<0.001, η_p_^2^=0.628, 1-β=1.000; Figure 5B). Within the older-aged group, TgAD rats showed larger peak amplitudes at higher intensities relative to WT (genotype: F_(1, 12)_=15.399, p=0.002, η_p_^2^=0.562, 1-β=0.949; intensity: F_(16, 192)_=52.101, p<0.001, η_p_^2^=0.813, 1-β=1.000; genotype × intensity: F_(16, 192)_=52.101, p=0.021, η_p_^2^=0.138, 1-β=0.952; Figure 5B).

#### Paired-pulse ratio

To determine whether the differences in the strength of basal transmission could be a consequence of differences in presynaptic release probability, we analyzed the paired-pulse ratio (PPR), an indirect measure of release probability (Dobrunz and Stevens, 1997). Analyzing the PPR at increasing stimulus intensities revealed that, irrespective of age and genotype, facilitation increased from 0.036 to 0.121 between 20 to 40µA, then decreased from 0.099 to 0.000 between 45 to 60µA before it reversed to depression from -0.035 to -0.114 between 65µA to 100µA (F_(16, 763)_=14.550, p<0.001, η_p_^2^=0.240, 1-β=1.000; see Table 3). No other effects were significant (Fs_(1–32, 46–736)_=0.298–1.750, ps=0.109–0.588). These findings indicate that presynaptic release probability does not explain the differences observed in the I/O curves and that the heightened transmission in the TgAD rats is unlikely to be caused by enhanced presynaptic glutamate release.

#### Hyperexcitability

To determine whether the heightened strength in basal transmission in young and aged TgAD rats is a consequence of hyperexcitability since there is no increase in presynaptic release probability, we analyzed the fEPSP traces using the coastline burst index (CBI). The value of the index (in arbitrary units) is associated with the intensity of the underlying activity in the fEPSP, (as previously reported in Korn et al., 1987; Stewart et al., 2017; Widman and McMahon, 2018). While CBIs did not differ by age (F_(2, 46)_=0.913, p=0.409, η_p_^2^=0.038, 1-β=0.198), all TgAD rats had a larger CBIs relative to all WT (+0.808; F_(1, 46)_=6.491, p=0.014, η_p_^2^=0.124, 1-β=0.703), and CBIs increased in all rats at larger stimulation intensities (F_(16, 736)_=41.445, p<0.001, η_p_^2^=0.474, 1-β=1.000), as expected when more axons are recruited at stronger stimulus intensities. Furthermore, there were significant genotype × intensity (F_(16, 736)_=4.175, p=0.020, η_p_^2^=0.083, 1-β=0.712) and trending age × intensity interactions (F_(32, 736)_=2.455, p=0.053, η_p_^2^=0.096, 1-β=0.670) with no other effects (Fs_(2–32, 46–736)_=0.040–0.405, p=0.799–0.960).

To better understand the reported interactions above, we tested if CBIs differed by genotype at each age. Indeed, within the young adult age, TgAD rats had larger CBIs at increasing intensities (genotype: F_(1, 15)_=5.202, p=0.038, η_p_^2^=0.257, 1-β=0.569; intensity: F_(16, 240)_=13.153, p<0.001, η_p_^2^=0.467, 1-β=1.000; genotype × intensity: (F_(16, 240)_=1.681, p=0.051, η_p_^2^=0.101, 1-β=0.919; Figure 5C). Within the middle-aged group, genotypes did not differ (Fs_(1–16, 19–304)_=1.059– 1.895, ps=0.177–0.316), but CBIs increased with increasing intensity in all rats as expected with greater recruitment of axons (F_(16, 304)_=19.393, p<0.001, η_p_^2^=0.505, 1-β=0.998; Figure 6A). Within the older-aged group, TgAD rats had larger CBIs with increasing intensity relative to WT (genotype: F_(1, 12)_=9.895, p=0.008, η_p_^2^=0.452, 1-β=0.823; intensity: F_(16, 192)_=49.361, p<0.001, η_p_^2^=0.804, 1-β=0.998; genotype × intensity: F_(16, 192)_=5.794, p<0.001, η_p_^2^=0.326, 1-β=1.000; Figure 5C).

**Figure 6.**
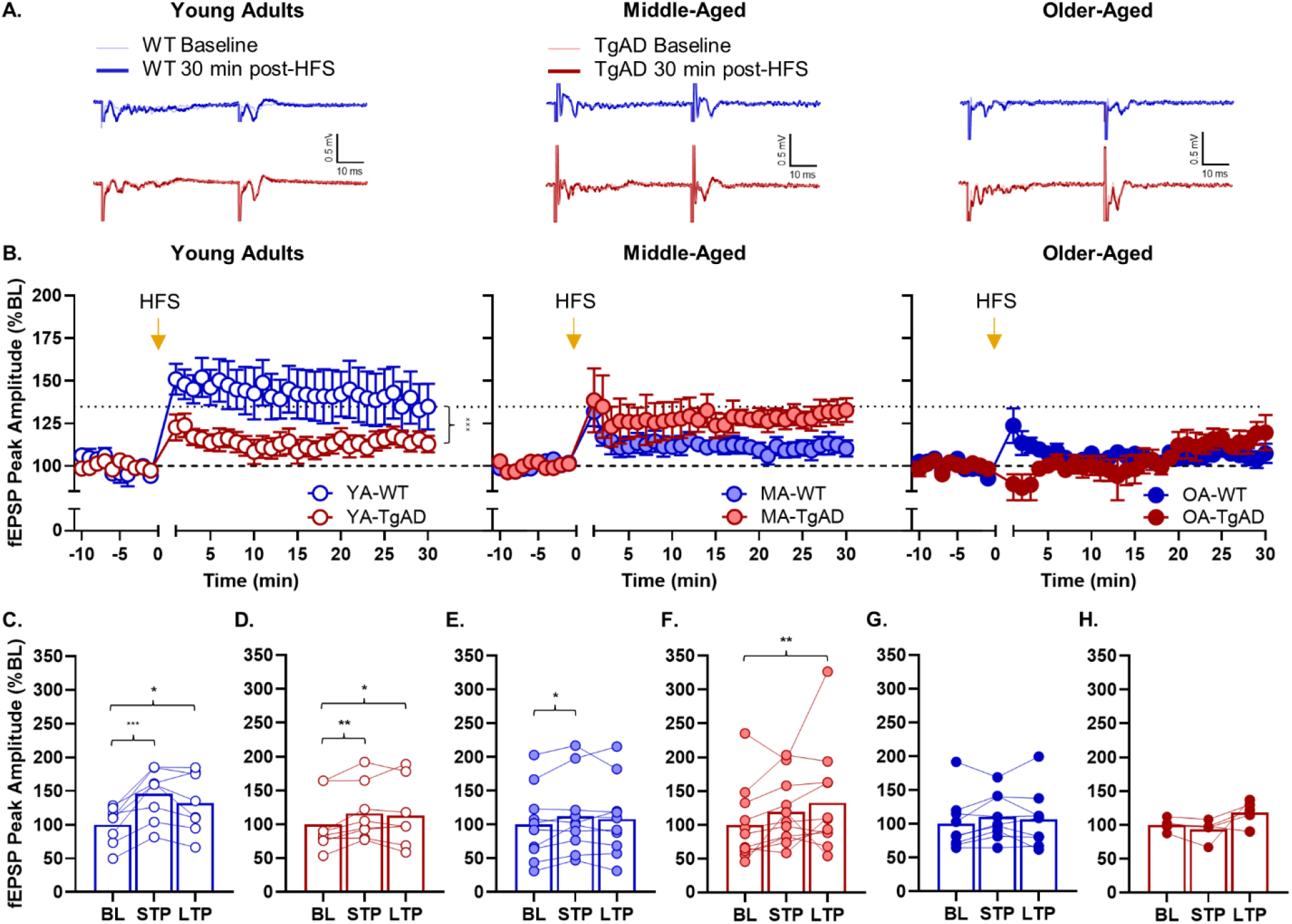
Effects of high-frequency stimulation on fEPSP peak amplitudes. **A)** Representative traces for each group. **B)** The fEPSP peak amplitude differences between baseline and post high-frequency stimulation (HFS) within each age group. There is a significant genotype by time interaction within the young adult group, whereas there are no genotypic differences within the other age groups. There was an overall main effect of age such that young adults had larger magnitude post-HFS peak amplitudes relative to middle- and older-aged groups. **C)** Short-term potentiation (STP) and long-term potentiation (LTP) relative to baseline (BL) within the young adult WT rats. There was significant STP and LTP relative to BL. **D)** STP and LTP relative to BL within the young adult TgAD rats. There was significant STP and LTP relative to BL. **E)** STP and LTP relative to BL within the middle-aged WT rats. There was significant STP but not LTP relative to BL. **F)** STP and LTP relative to BL within the middle-aged TgAD rats. While there was not significant STP, there was significant LTP relative to BL. **G)** STP and LTP relative to BL within the older-aged WT rats. There was no STP or LTP relative to BL. **H)** STP and LTP relative to BL within the older-aged TgAD rats. There was no STP or LTP relative to BL. Data are plotted as the means and SEM and fEPSP peak amplitudes as percent of baseline. *p<0.05, **p<0.01, ***p<0.001 for paired-samples t-tests; ×××p<0.001 for genotype × time interactions. Dotted black line represents mean of YA-WT at 30 minutes post-HFS. Dashed black line represents baseline.

To confirm genotypic differences in baseline synaptic strength and hyperexcitability were not explained by differences in the pre-stimulus period, we analyzed CBIs during the pre-stimulus period. Although there were no effects of intensity, genotype, or interactions (Fs_(1–32, 46–736)_=0.039– 0.946, ps=0.489–0.951), there was a small magnitude effect of age (F_(2, 46)_=5.153, p=0.010, η_p_^2^=0.183, 1-β=0.801) such that middle-aged rats showed smaller pre-stimulus CBIs relative to young adult (−0.189, p=0.017) and older-aged rats (−0.238, p=0.006; data not shown). However, a small magnitude decrease in the middle-aged CBI pre-stimulus period does not explain the observed increase in CBIs driven by age and genotype.

Taken together, these data suggest the BA of TgAD rats is overall hyperexcitable. The null effects of genotype on BA activity in middle-aged rats are consistent with the ephemeral compensation in acute fear memory extinction observed in the middle-aged TgAD rats. These data suggest the BA of TgAD rats is hyperexcitable during young adulthood, undergoes a compensatory mechanism during middle-age but reverts to a hyperexcitable state in older-aged rats.

#### Long-term Potentiation

Given the changes in basal synaptic transmission, we next asked if synaptic plasticity was inhibited in TgAD and WT rats across the lifespan. In Figure 6A, representative traces show peak amplitudes during baseline and post-high frequency stimulation (HFS) for each group. As reported previously (Zeng et al., 2012; Zhan et al., 2018), the maximum peak amplitude decreases with age (F_(2, 45)_=4.036, p=0.024, η_p_^2^=0.152, 1-β=0.691) such that older-aged rats showed smaller peak amplitudes across all post-HFS time points relative to young adult (−16.394, p=0.007) and middle-aged (−11.215, p=0.049) rats (Figure 6B). While there was no main effect of genotype (F_(1, 45)_=0.571, p=0.454), there was an expected effect of time (F_(39, 1755)_=12.454, p<0.001, η_p_^2^=0.217, 1-β=1.000) indicating expected increases in peak amplitudes post-tetanus across all groups. Importantly, age interacted with all factors whereas genotype did not interact with time (age × genotype: F_(2, 45)_=4.766, p=0.013, η_p_^2^=0.175, 1-β=0.766; age × time: F_(78, 1755)_=2.101, p=0.029, η_p_^2^=0.085, 1-β=0.874; age × genotype × time: F_(78, 1755)_=2.079, p=0.031, η_p_^2^=0.085, 1-β=0.870; genotype × time: F_(39, 1755)_=1.396, p=0.231).

We then asked if there were general magnitude differences between genotype groups at each age. Within the young adults, peak amplitudes at all time points after HFS were larger in WT relative to TgAD rats (genotype: F_(1, 14)_=3.615, p=0.078; time: F_(39, 546)_=9.161, p<0.001, η_p_^2^=0.396, 1-β=1.000; genotype × time: F_(39, 546)_=2.399, p<0.001, η_p_^2^=0.146, 1-β=1.000; Figure 7B). Within the middle-aged group, though there were numerical differences, the magnitude of peak amplitudes were not statistically different between WT and TgAD rats (Fs_(1–39, 19–741)_=1.042-3.316, ps=0.084–0.402). However, peak amplitudes were greater at all timepoints after HFS relative to baseline in all middle-aged rats (time: F_(39, 741)_=6.088, p<0.001, η_p_^2^=0.243, 1-β=1.000). Within the older-aged group, TgAD rats showed decreased peak amplitudes only within the first 3 minutes post-HFS relative to WT rats (genotype: F_(1, 12)_=0.406, p=0.536; time: F_(39, 468)_=2.181, p<0.001, η_p_^2^=0.154, 1-β=1.000; genotype × time: F_(39, 468)_=2.668, p<0.001, η_p_^2^=0.182, 1-β=1.000; Figure 7B). Finally, these effects were not explained by baseline differences in peak amplitudes (age × genotype × time ANOVA: Fs_(1–39, 19–741)_=3.363–1.410, ps=0.197–0.830).

**Figure 7.**
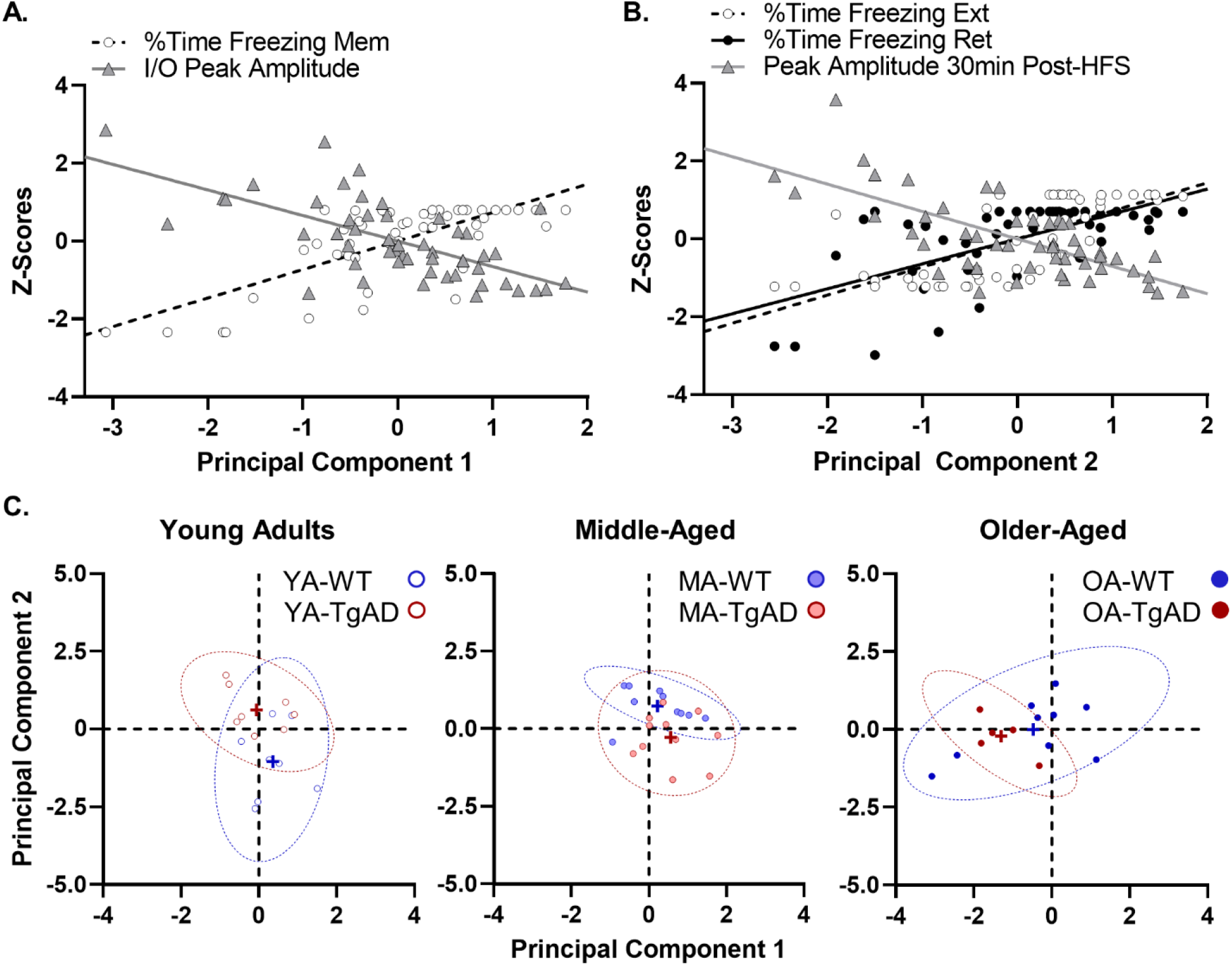
Principal component analysis on measures of CS extinction and BA synaptic function. **A)** Significant correlations with component 1. In rats with large I/O peak amplitudes, there was also less freezing during the memory probe test. **B)** significant correlations with component 2. In rats with impaired LTP, there was also impaired extinction and extinction memory retrieval. **C)** Group differences in the contributions to each component resulted in unique clustering. Component 1 segregated rats by age such that all older-aged rats significantly clustered more negatively than young or middle-aged rats indicating the rats at the tail end of the correlations with component 1 displayed in panel A were older-aged rats. In contrast, component 2 segregated groups by age and genotype such that young adult WT rats clustered more negatively relative to middle- and older-aged WT counterparts and their young TgAD counterparts. This confirmed that the rats at the negative tail of the correlations with component 2 displayed in panel B were young adult WT rats. Additionally, middle-aged TgAD rats also clustered more negatively than their young TgAD or middle-aged WT counterparts indicating the rats more positively adjacent to young WT in the panel B correlations were middle-aged TgAD rats. While the middle-aged TgAD rats had a minor young-like phenotype, the more positive shift was likely due to the impairment in extinction memory retrieval. In panel C, crosses represent group means, and dotted ellipses represent the 95% confidence intervals.

In addition to testing group differences, we wanted to confirm the degree to which WT and TgAD rats across the lifespan were capable of STP and LTP by using paired-samples t-tests. Young adult WT rats had larger peak amplitudes within the first 5 minutes (t_(7)_=5.525, p<0.001, *d*=1.953) and also at 30 minutes (t_(7)_=2.414, p=0.047, *d*=0.853) post-HFS (Figure 7C). Young adult TgAD rats had larger peak amplitudes within the first 5 minutes (t_(7)_=3.761, p=0.007, *d*=1.330) and also at 30 minutes (t_(7)_=2.929, p=0.022, *d*=1.036) post-HFS (Figure 7D). Middle-aged WT rats showed larger peak amplitudes within the first 5 minutes (t_(9)_=2.926, p=0.017, *d*=0.925) but not at 30 minutes (t_(9)_=1.415, p=0.191) post-HFS (Figure 7E). Middle-aged TgAD rats did not show a difference in peak amplitudes within the first 5 minutes (t_(10)_=2.117, p=0.060), however, peak amplitudes were larger at 30 minutes (t_(10)_=4.023, p=0.002) post-HFS (Figure 7F). In contrast to young adult and middle-aged rats, there were no differences between baseline and post-HFS peak amplitudes in older-aged WT rats (ts_(10)_=1.300-1.978, ps=0.083-0.230; Figure 7G) and TgAD (ts_(4)_=-2.041–1.853, ps=0.111–0.138; Figure 7H). Taken together, these data suggest the attenuated LTP in young adult TgAD rats undergoes compensation during middle-age and is not sustain into older-age, whereas in WT rats aging significantly impairs BA LTP.

### Associations between behavior and synaptic physiology

The results above demonstrate larger I/O peak amplitudes and smaller magnitude or impaired LTP in groups with impaired extinction or retrieval and smaller I/O peak amplitudes with larger magnitude LTP in groups with intact extinction. As such, it is possible measures of BA synaptic physiology can predict fear memory expression during phases of extinction testing. Therefore, a PCA was used to determine if associations between measures of synaptic physiology and measures of extinction segregated groups based on individual variability. The final PCA used n=51 rats (note that due to a technical issue, n=1 young adult WT rat was removed). The Kaiser-Meyer-Olkin (KMO) measure of sampling adequacy was 0.630, and Bartlett’s test of sphericity was significant (≈χ^2^_(10)_=27.294, p=0.002) indicating a correlation between loaded variables. Item communalities were moderate to high (ranging from 0.577 to 0.668) for all items except the I/O peak amplitudes which was low (0.431).

A model with 2 components (Eigenvalues >1) explained 59.15% of the variance in the data. Component 1, which explained 38.18% of the variance, loaded positively with freezing during the CS memory probe trial (r=+0.731), but loaded negatively with the magnitude of peak amplitudes across all stimulation intensities (r=-0.656). These data indicate that rats with less fear memory expression during the CS memory probe had a more hyperexcitable BA (Figure 7A). In contrast, component 2, which explained 20.97% of the variance, loaded positively with freezing during extinction (r=+0.722) and extinction memory retrieval (r=+0.640), but loaded negatively with the magnitude of peak amplitudes at 30 minutes post-HFS (r=-0.704). These data indicate that rats with impaired extinction and extinction memory retrieval also had impaired BA LTP (Figure 7B).

We then tested the possibility that the individual variability within each age and genotype group loaded in a manner that contributed to components uniquely. Figure 7C shows component 1 significantly segregated groups by age (F_(2, 45)_=8.899, p<0.001, η_p_^2^=0.283, 1-β=0.963) such that older-aged rats clustered more negatively relative to young adults (−1.037, p=0.003) and middle-aged rats (−1.277, p<0.001), whereas genotype had no effect nor did it interact with age (Fs_(1–2, 45)_=1.446–1.927, ps=0.157–0.236). This indicated that older-aged rats demonstrated the highest I/O peak amplitudes corresponding to the lowest freezing during the memory probe trial (Mem).

In other words, older-aged rats clustered toward the negative tail of the correlations with component 1. Figure 7C also shows component 2 significantly segregated age groups by genotype (F_(2, 45)_=11.444, p<0.001, η_p_^2^>=0.337, 1-β=0.990) with no main effects (Fs_(1–2, 45)_=0.427– 1.444, ps=0.247–0.517). Specifically, young adult WT rats clustered more negatively relative to middle-aged (−1.776, p<0.001) and older-aged WT (−1.042, p=0.014) rats and their young adult TgAD counterparts (−1.664, p<0.001). This indicated that young adult WT rats as a group showed the best fear extinction, extinction memory retrieval, and largest peak amplitudes at 30 min post-HFS. Indeed, the young adult WT rats clustered toward the negative tail of the correlations with component 2. Interestingly, middle-aged TgAD rats also clustered more negatively relative to young adult TgAD rats (−0.874, p=0.031) and their middle-aged WT counterparts (−0.986, p=0.010). Although they were adjacent to the young adult WT rats, the middle-aged TgAD rats did not occupy the same portion of the correlation that young adult WT rats occupied, as they were impaired in extinction memory retrieval despite having intact extinction and LTP. Together, these data suggest two possibilities. First, irrespective of genotype, BA hyperexcitability in old age may impact the degree to which fear memory is expressed, though not necessarily fear memory per se. Secondly, BA LTP facilitates the degree to which fear memory extinction and extinction memory retrieval occurs. Importantly, these results are consistent with the age × genotype interactions in the behavioral and electrophysiological measures noted above and further agree with the compensatory mechanisms observed in the middle-aged TgAD rats that do not extend into old age.

## Discussion

In summary, non-pathological aging and AD impair extinction memory and generalize contextual fear memory. While aging impairs LTP overall, it occurs later in TgAD rats, and the BA is hyperexcitable across the lifespan in TgAD but not WT rats. Importantly, these findings are not explained by differences in shock perception, environmental habituation (Oler and Markus, 1998; Moyer and Brown, 2006; Luo et al., 2015; Zhan et al., 2018), or the pre-stimulus period during synaptic recordings.

Consistent with no genotypic difference in fear acquisition in young and middle-aged rats, we recently reported no differences in contextual fear acquisition in a cohort of 10–13-mo-old WT and TgAD rats given a series of uncued foot shocks (Goodman et al., 2021). Aging, however, attenuated acquisition in middle-aged rats, spared acquisition in older-aged WT rats, and impaired acquisition in older-aged TgAD rats, consistent with studies showing aging attenuates, but doesn’t grossly impair, acquisition (Villarreal et al., 2004; Kaczorowski et al., 2012). It is possible that all middle-aged rats and older-aged TgAD rats would show enhanced fear acquisition given a larger number of trials or shorter delays between CS onset and US delivery (Maren, 1999; McEchron et al., 2004; Villarreal et al., 2004; Moyer and Brown, 2006; Detert et al., 2008). Our results in older-aged TgAD rats are consistent with impaired fear memory acquisition in Alzheimer’s patients (Hamann et al., 2002; Hoefer et al., 2008).

Rats at all ages showed increased fear memory expression during the memory probe trial, suggesting there was a 24-hr fear memory incubation period (Herry et al., 2008). The fact that acute acquisition deficits were overcome suggests some degree of intact fear memory consolidation and retrieval. The BA has two populations of glutamatergic neurons, known as “fear” and “extinction” neurons (Herry et al., 2008; Amano et al., 2011; Pare and Duvarci, 2012). One hypothesis for the emergence of memory in the older-aged TgAD rats is that hyperexcitable BA “fear” neurons facilitated consolidation and retrieval. Indeed, older-aged rats showed both greater hyperexcitability and less fear expression during the memory probe trial.

Greater fear expression in young TgAD rats during the probe trial is consistent with enhanced fear to cued foot shocks in AD mouse models (España et al., 2010). Furthermore, impaired extinction in young TgAD rats is consistent with deficits in 4.5-month-old APP^swe^/PS1^ΔE9^ (Bonardi et al., 2011), and TASTPM mice (Rattray et al., 2009). Fear extinction requires that retrieved memories be labile and conducive to inhibition and updating prior to reconsolidation (Kida, 2019). Moreover, BA “extinction” neurons may need to be disinhibited from upstream hyperexcitable “fear” neurons via an intermediary inhibitory circuit to permit extinction (Pare and Duvarci, 2012). Therefore, the hyperexcitability at BA synapses in young TgAD rats suggests an insurmountable drive on “fear” neurons, rendering the memory resistant to a labile state. Moreover, attenuated LTP in young TgAD rats suggests an inflexible circuit and further undermines the reinforcement of extinction. These data further support the evidence that emotional memory deficits are an early sign of AD (Hamann et al., 2002; Hoefer et al., 2008) and are consistent with known amygdala-related deficits in AD (Wright et al., 2007).

Though unexpected, enhanced extinction in middle-aged TgAD rats is consistent with our previous study of no extinction differences between 10–13-month-old WT and TgAD rats (Goodman et al., 2021) and another reporting 3×Tg-AD mice were better than WT during extinction (Pietropaolo et al., 2008). As detailed above, it is unlikely that diminished salience to the US explains spared extinction, though circuit-level compensation could offer one explanation. Despite hyperexcitability in the middle-aged TgAD BA, synaptic strength and LTP mimic young adult WT. This young-WT-like phenotype may facilitate acute extinction encoding but not extinction memory. Though aging is not usually emphasized in rodent models of AD, morphological and synaptic function changes occur in the absence of cell death in the BLA of middle-aged APP^swe^/PS1^ΔE9^ mice (Knafo et al., 2009), suggesting that middle age is accompanied by a local network restructuring prior to neuronal loss that facilitates an ephemeral compensation in extinction and BA function.

This compensation, however, does not extend into older age. The persistence of BA hyperexcitability in middle-age TgAD rats may set the stage for a vulnerable environment negatively impacted by aging. Consistent with our findings, fear extinction is progressively worse in AD relative to MCI patients and healthy controls (Nasrouei et al., 2020), and AD patients show heighted amygdala responses to negative stimuli (Wright et al., 2007). Young adult and middle-aged TgAD rats may be displaying memory deficits akin to MCI, and by old age, those deficits are compounded into clear impairments in emotional memory driven by underlying pathological changes resulting in hyperactive and rigid BA synapses.

One strength of our study is addressing aging within the context of AD. Indeed, our results provide further evidence that non-pathological aging is distinct from pathological aging, as extinction and synaptic deficits in WT rats across the lifespan do not mirror those in TgAD rats. A second strength of our study is the inclusion of a middle-aged cohort to better define the trajectory of outcomes across the lifespan. Similarly, another study showed impaired extinction in middle-aged WT rats (Kaczorowski et al., 2012), and some degree of extinction in older-aged WT rats, albeit attenuated in magnitude (Moyer and Brown, 2006; Kaczorowski et al., 2012). However, Villarreal et al., 2004 reported impaired extinction in 22-month-old rats. While impaired LTP in the BA of middle-aged WT rats is novel, consistent with our own findings, older age is accompanied by impaired LTP (Zeng et al., 2012; Zhan et al., 2018). These results were not surprising given a wide body of literature showing older adults and rats depend on differential neural recruitment to achieve equal performance (Samanez-Larkin et al., 2011; Antonenko and Flöel, 2014; Lighthall et al., 2014; Tomás Pereira et al., 2015; Wang et al., 2015; Hernandez et al., 2019). Notably, these studies did not incorporate a middle-aged group to determine whether age differences are progressive, abrupt, or dynamic. Older-aged WT rats may require extra-amygdalar circuit recruitment during extinction not necessary in middle-aged WT rats.

Aging and AD significantly impaired extinction memory retrieval. Only young adult WT rats retrieved extinction memory. Fear memories that undergo extinction are reconsolidated as new memories while the original memory remains with less saliency (Bouton and King, 1983; Barad et al., 2006). In rats showing acute extinction, it is difficult to determine if impaired retrieval was due to poor reconsolidation or an impairment in retrieval itself. Nonetheless, both mnemonic mechanisms are supported by the BLA, and the BA nucleus specifically (Maren, 2001; Kochli et al., 2015), and as such retrieval in aging and AD may rely on an intact BA for extinction encoding and reconsolidation. It is possible synaptic compensation in middle-aged TgAD rats affords the encoding of acute extinction memory that does not appropriately reconsolidate, whereas additional circuitry recruitment facilitates extinction encoding but not reconsolidation in older-aged WT rats. The associations between extinction retrieval and synaptic function in aged WT and all TgAD rats deviate from young WT suggesting any combination of synaptic deficits results in long-term extinction deficits. Although the BLA is a foci for fear memory encoding, consolidation, and storage, other neural circuitry can compensate when BLA deficits are present (Cahill et al., 2000; Maren, 2001; Kochli et al., 2015), particularly in aging (Hernandez et al., 2019).

Our findings suggest a maladaptive expression of fear despite clear discrimination between safe and unsafe contexts in older-aged WT rats. Consistent with the current study, 23-month-old rats demonstrated comparable fear expression during contextual renewal testing relative to young adults suggesting intact contextual discrimination and memory (Oler and Markus, 1998). In all other groups, however, there was progressive increases in contextual fear memory expression, suggesting a maladaptive fear response similarly observed in neuropsychiatric disorders like posttraumatic stress disorder (Garfinkel et al., 2014; Huckleberry et al., 2016). Although the hippocampus is critical in contextual fear memory consolidation (Phillips and LeDoux, 1992), there is evidence of hippocampal-independent mechanisms of consolidation (Wiltgen et al., 2006; Kochli et al., 2015). The context deficits in TgAD rats are similar throughout all stages of life, and it is reasonable to suggest the contribution to extinction impairments and contextual generalization is shared between persistent BA hyperactivity and known hippocampal deficits in this rat model (Smith and McMahon, 2018; Goodman et al., 2021). In non-pathological aging, contextual fear memory deficits may also be explained by BA and hippocampal dysfunction given the wide body of literature showing age-related impairments to hippocampal function (Comery et al., 2005; Ohno, 2009; Zeng et al., 2012; Fjell et al., 2014; Zhan et al., 2018; Burke and Foster, 2019).

## Conclusion

These results emphasize the unique trajectories aging takes in the presence and absence of disease. As AD risk significantly increases with age, and potential compensatory mechanisms may confound interpretations of data that do not account for age, more studies would benefit from incorporating age as a factor when using rodent models of AD.

## Acknowledgements

The work was supported by NIH/NICHD 2T32HD071866-06 to CMH and ARH; NIH/NIA R01AG066489 to LLM.

## Abbreviations

(BLA): basolateral amygdala
(BA): basal amygdaloid nucleus
(fEPSP): field excitatory postsynaptic potential
(LTP): long-term potentiation

## References

Amano T, Duvarci S, Popa D, Paré D (2011) The fear circuit revisited: contributions of the basal amygdala nuclei to conditioned fear. J Neurosci 31:15481–15489.

Amorapanth P, LeDoux JE, Nader K (2000) Different lateral amygdala outputs mediate reactions and actions elicited by a fear-arousing stimulus. Nat Neurosci 3:74–79.

Antonenko D, Flöel A (2014) Healthy aging by staying selectively connected: a mini-review. Gerontology 60:3–9.

Barad M, Gean P-W, Lutz B (2006) The role of the amygdala in the extinction of conditioned fear. Biol Psychiatry 60:322–328.

Beaudreau SA, O’Hara R (2008) Late-life anxiety and cognitive impairment: a review. Am J Geriatr Psychiatry 16:790–803.

Belova MA, Paton JJ, Salzman CD (2008) Moment-to-moment tracking of state value in the amygdala. J Neurosci 28:10023–10030.

Beyeler A, Namburi P, Glober GF, Simonnet C, Calhoon GG, Conyers GF, Luck R, Wildes CP, Tye KM (2016) Divergent Routing of Positive and Negative Information from the Amygdala during Memory Retrieval. Neuron 90:348–361.

Bonardi C, de Pulford F, Jennings D, Pardon M-C (2011) A detailed analysis of the early context extinction deficits seen in APPswe/PS1dE9 female mice and their relevance to preclinical Alzheimer’s disease. Behav Brain Res 222:89–97.

Bouton ME, King DA (1983) Contextual control of the extinction of conditioned fear: tests for the associative value of the context. J Exp Psychol Anim Behav Process 9:248–265.

Buckner RL (2004) Memory and executive function in aging and AD: multiple factors that cause decline and reserve factors that compensate. Neuron 44:195–208.

Burke SL, Cadet T, Alcide A, O’Driscoll J, Maramaldi P (2018) Psychosocial risk factors and Alzheimer’s disease: the associative effect of depression, sleep disturbance, and anxiety. Aging Ment Health 22:1577–1584.

Burke SN, Foster TC (2019) Animal models of cognitive aging and circuit-specific vulnerability. Handb Clin Neurol 167:19–36.

Cahill L, Vazdarjanova A, Setlow B (2000) The basolateral amygdala complex is involved with, but is not necessary for, rapid acquisition of Pavlovian ‘fear conditioning. European Journal of Neuroscience 12:3044–3050.

Cohen RM, Rezai-Zadeh K, Weitz TM, Rentsendorj A, Gate D, Spivak I, Bholat Y, Vasilevko V, Glabe CG, Breunig JJ, Rakic P, Davtyan H, Agadjanyan MG, Kepe V, Barrio JR, Bannykh S, Szekely CA, Pechnick RN, Town T (2013) A transgenic Alzheimer rat with plaques, tau pathology, behavioral impairment, oligomeric aβ, and frank neuronal loss. J Neurosci 33:6245–6256.

Comery TA, Martone RL, Aschmies S, Atchison KP, Diamantidis G, Gong X, Zhou H, Kreft AF, Pangalos MN, Sonnenberg-Reines J, Jacobsen JS, Marquis KL (2005) Acute gamma-secretase inhibition improves contextual fear conditioning in the Tg2576 mouse model of Alzheimer’s disease. J Neurosci 25:8898–8902.

Detert JA, Kampa ND, Moyer JR (2008) Differential effects of training intertrial interval on acquisition of trace and long-delay fear conditioning in rats. Behav Neurosci 122:1318–1327.

DiStefano C, Zhu M, Mîndrilã D (2009) Understanding and Using Factor Scores: Considerations for the Applied Researcher. University of Massachusetts Amherst.

Dobrunz LE, Stevens CF (1997) Heterogeneity of release probability, facilitation, and depletion at central synapses. Neuron 18:995–1008.

España J, Giménez-Llort L, Valero J, Miñano A, Rábano A, Rodriguez-Alvarez J, LaFerla FM, Saura CA (2010) Intraneuronal beta-amyloid accumulation in the amygdala enhances fear and anxiety in Alzheimer’s disease transgenic mice. Biol Psychiatry 67:513–521.

Fjell AM, McEvoy L, Holland D, Dale AM, Walhovd KB, Alzheimer’s Disease Neuroimaging Initiative (2014) What is normal in normal aging? Effects of aging, amyloid and Alzheimer’s disease on the cerebral cortex and the hippocampus. Prog Neurobiol 117:20–40.

Floyd M, Rice J, Black SR (2002) Recurrence of Posttraumatic Stress Disorder in Late Life: A Cognitive Aging Perspective. Journal of Clinical Geropsychology.

Garavan H, Pendergrass JC, Ross TJ, Stein EA, Risinger RC (2001) Amygdala response to both positively and negatively valenced stimuli. Neuroreport 12:2779–2783.

Garfinkel SN, Abelson JL, King AP, Sripada RK, Wang X, Gaines LM, Liberzon I (2014) Impaired contextual modulation of memories in PTSD: an fMRI and psychophysiological study of extinction retention and fear renewal. J Neurosci 34:13435–13443.

Goodman AM, Langner BM, Jackson N, Alex C, McMahon LL (2021) Heightened Hippocampal β-Adrenergic Receptor Function Drives Synaptic Potentiation and Supports Learning and Memory in the TgF344-AD Rat Model during Prodromal Alzheimer’s Disease. J Neurosci 41:5747–5761.

Green E, Fairchild JK, Kinoshita LM, Noda A, Yesavage J (2016) Effects of posttraumatic stress disorder and metabolic syndrome on cognitive aging in veterans. Gerontologist 56:72–81.

Hamann S, Monarch ES, Goldstein FC (2002) Impaired fear conditioning in Alzheimer’s disease. Neuropsychologia 40:1187–1195.

Hernandez CM, Orsini C, Wheeler A-R, Ten Eyck TW, Betzhold SM, Labiste CC, Wright NG, Setlow B, Bizon JL (2020) Testicular hormones mediate robust sex differences in impulsive choice in rats. Elife 9.

Hernandez CM, Orsini CA, Labiste CC, Wheeler A-R, Ten Eyck TW, Bruner MM, Sahagian TJ, Harden SW, Frazier CJ, Setlow B, Bizon JL (2019) Optogenetic dissection of basolateral amygdala contributions to intertemporal choice in young and aged rats. Elife 8.

Herry C, Ciocchi S, Senn V, Demmou L, Müller C, Lüthi A (2008) Switching on and off fear by distinct neuronal circuits. Nature 454:600–606.

Hoefer M, Allison SC, Schauer GF, Neuhaus JM, Hall J, Dang JN, Weiner MW, Miller BL, Rosen HJ (2008) Fear conditioning in frontotemporal lobar degeneration and Alzheimer’s disease. Brain 131:1646–1657.

Huang Y-Y, Kandel ER (2007) 5-Hydroxytryptamine induces a protein kinase A/mitogen-activated protein kinase-mediated and macromolecular synthesis-dependent late phase of long-term potentiation in the amygdala. J Neurosci 27:3111–3119.

Huckleberry KA, Ferguson LB, Drew MR (2016) Behavioral mechanisms of context fear generalization in mice. Learn Mem 23:703–709.

Humeau Y, Reisel D, Johnson AW, Borchardt T, Jensen V, Gebhardt C, Bosch V, Gass P, Bannerman DM, Good MA, Hvalby Ø, Sprengel R, Lüthi A (2007) A pathway-specific function for different AMPA receptor subunits in amygdala long-term potentiation and fear conditioning. J Neurosci 27:10947–10956.

Kaczorowski CC, Davis SJ, Moyer JR (2012) Aging redistributes medial prefrontal neuronal excitability and impedes extinction of trace fear conditioning. Neurobiol Aging 33:1744–1757.

Kida S (2019) Reconsolidation/destabilization, extinction and forgetting of fear memory as therapeutic targets for PTSD. Psychopharmacology 236:49–57.

Knafo S, Venero C, Merino-Serrais P, Fernaud-Espinosa I, Gonzalez-Soriano J, Ferrer I, Santpere G, DeFelipe J (2009) Morphological alterations to neurons of the amygdala and impaired fear conditioning in a transgenic mouse model of Alzheimer’s disease. J Pathol 219:41–51.

Kochli DE, Thompson EC, Fricke EA, Postle AF, Quinn JJ (2015) The amygdala is critical for trace, delay, and contextual fear conditioning. Learn Mem 22:92–100.

Korn SJ, Giacchino JL, Chamberlin NL, Dingledine R (1987) Epileptiform burst activity induced by potassium in the hippocampus and its regulation by GABA-mediated inhibition. J Neurophysiol 57:325–340.

LeDoux JE, Cicchetti P, Xagoraris A, Romanski LM (1990) The lateral amygdaloid nucleus: sensory interface of the amygdala in fear conditioning. J Neurosci 10:1062–1069.

Lighthall NR, Huettel SA, Cabeza R (2014) Functional compensation in the ventromedial prefrontal cortex improves memory-dependent decisions in older adults. J Neurosci 34:15648–15657.

Liley AE, Gabriel DBK, Sable HJ, Simon NW (2019) Sex differences and effects of predictive cues on delayed punishment discounting. eNeuro 6.

Luo Y, Zhou J, Li M-X, Wu P-F, Hu Z-L, Ni L, Jin Y, Chen J-G, Wang F (2015) Reversal of aging-related emotional memory deficits by norepinephrine via regulating the stability of surface AMPA receptors. Aging Cell 14:170–179.

Maren S (1999) Neurotoxic basolateral amygdala lesions impair learning and memory but not the performance of conditional fear in rats. J Neurosci 19:8696–8703.

Maren S (2001) Is there savings for pavlovian fear conditioning after neurotoxic basolateral amygdala lesions in rats? Neurobiol Learn Mem 76:268–283.

Maren S, De Oca B, Fanselow MS (1994) Sex differences in hippocampal long-term potentiation (LTP) and Pavlovian fear conditioning in rats: positive correlation between LTP and contextual learning. Brain Res 661:25–34.

Maren S, Quirk GJ (2004) Neuronal signalling of fear memory. Nat Rev Neurosci 5:844–852.

Mcdonald AJ, Mascagni F, Guo L (1996) Projections of the medial and lateral prefrontal cortices to the amygdala: a Phaseolus vulgaris leucoagglutinin study in the rat. Neuroscience 71:55–75.

McDonald AJ, Mott DD (2017) Functional neuroanatomy of amygdalohippocampal interconnections and their role in learning and memory. J Neurosci Res 95:797–820.

McEchron MD, Cheng AY, Gilmartin MR (2004) Trace fear conditioning is reduced in the aging rat. Neurobiol Learn Mem 82:71–76.

McKhann GM, Albert MS, Grossman M, Miller B, Dickson D, Trojanowski JQ, Work Group on Frontotemporal Dementia and Pick’s Disease (2001) Clinical and pathological diagnosis of frontotemporal dementia: report of the Work Group on Frontotemporal Dementia and Pick’s Disease. Arch Neurol 58:1803–1809.

Miettunen J, Veijola J, Lauronen E, Kantojärvi L, Joukamaa M (2007) Sex differences in Cloninger’s temperament dimensions--a meta-analysis. Compr Psychiatry 48:161–169.

Mohamed AZ, Cumming P, Götz J, Nasrallah F, Department of Defense Alzheimer’s Disease Neuroimaging Initiative (2019) Tauopathy in veterans with long-term posttraumatic stress disorder and traumatic brain injury. Eur J Nucl Med Mol Imaging 46:1139–1151.

Moyer JR, Brown TH (2006) Impaired trace and contextual fear conditioning in aged rats. Behav Neurosci 120:612–624.

Nasrouei S, Rattel JA, Liedlgruber M, Marksteiner J, Wilhelm FH (2020) Fear acquisition and extinction deficits in amnestic mild cognitive impairment and early Alzheimer’s disease. Neurobiol Aging 87:26–34.

National Institute on Aging (2019) What Causes Alzheimer’s Disease? Causes of Alzheimer’s Disease Available at: https://www.nia.nih.gov/health/what-causes-alzheimers-disease [Accessed July 13, 2021].

Neary D, Snowden JS, Gustafson L, Passant U, Stuss D, Black S, Freedman M, Kertesz A, Robert PH, Albert M, Boone K, Miller BL, Cummings J, Benson DF (1998) Frontotemporal lobar degeneration: a consensus on clinical diagnostic criteria. Neurology 51:1546–1554.

Ohno M (2009) Failures to reconsolidate memory in a mouse model of Alzheimer’s disease. Neurobiol Learn Mem 92:455–459.

Oler JA, Markus EJ (1998) Age-related deficits on the radial maze and in fear conditioning: Hippocampal processing and consolidation. Hippocampus 8:402–415.

Orsini CA, Willis ML, Gilbert RJ, Bizon JL, Setlow B (2016) Sex differences in a rat model of risky decision making. Behav Neurosci 130:50–61.

Pare D, Duvarci S (2012) Amygdala microcircuits mediating fear expression and extinction. Curr Opin Neurobiol 22:717–723.

Pare WP (1965) Stress and consummatory behavior in the albino rat. Psychol Rep 16:399–405.

Peters J, Kalivas PW, Quirk GJ (2009) Extinction circuits for fear and addiction overlap in prefrontal cortex. Learn Mem 16:279–288.

Phillips RG, LeDoux JE (1992) Differential contribution of amygdala and hippocampus to cued and contextual fear conditioning. Behav Neurosci 106:274–285.

Pietropaolo S, Feldon J, Yee BK (2008) Age-dependent phenotypic characteristics of a triple transgenic mouse model of Alzheimer disease. Behav Neurosci 122:733–747.

Pryce CR, Lehmann J, Feldon J (1999) Effect of sex on fear conditioning is similar for context and discrete CS in Wistar, Lewis and Fischer rat strains. Pharmacol Biochem Behav 64:753–759.

Qureshi SU, Kimbrell T, Pyne JM, Magruder KM, Hudson TJ, Petersen NJ, Yu H-J, Schulz PE, Kunik ME (2010) Greater prevalence and incidence of dementia in older veterans with posttraumatic stress disorder. J Am Geriatr Soc 58:1627–1633.

Rattray I, Scullion GA, Soulby A, Kendall DA, Pardon M-C (2009) The occurrence of a deficit in contextual fear extinction in adult amyloid-over-expressing TASTPM mice is independent of the strength of conditioning but can be prevented by mild novel cage stress. Behav Brain Res 200:83–90.

Romanski LM, Clugnet MC, Bordi F, LeDoux JE (1993) Somatosensory and auditory convergence in the lateral nucleus of the amygdala. Behav Neurosci 107:444–450.

Samanez-Larkin GR, Mata R, Radu PT, Ballard IC, Carstensen LL, McClure SM (2011) Age Differences in Striatal Delay Sensitivity during Intertemporal Choice in Healthy Adults. Front Neurosci 5:126.

Shin LM, Rauch SL, Pitman RK (2006) Amygdala, medial prefrontal cortex, and hippocampal function in PTSD. Ann N Y Acad Sci 1071:67–79.

Sierra-Mercado D, Padilla-Coreano N, Quirk GJ (2011) Dissociable roles of prelimbic and infralimbic cortices, ventral hippocampus, and basolateral amygdala in the expression and extinction of conditioned fear. Neuropsychopharmacology 36:529–538.

Smith LA, McMahon LL (2018) Deficits in synaptic function occur at medial perforant path-dentate granule cell synapses prior to Schaffer collateral-CA1 pyramidal cell synapses in the novel TgF344-Alzheimer’s Disease Rat Model. Neurobiol Dis 110:166–179.

Stewart LT, Khan AU, Wang K, Pizarro D, Pati S, Buckingham SC, Olsen ML, Chatham JC, McMahon LL (2017) Acute Increases in Protein O-GlcNAcylation Dampen Epileptiform Activity in Hippocampus. J Neurosci 37:8207–8215.

Stover KR, Campbell MA, Van Winssen CM, Brown RE (2015) Early detection of cognitive deficits in the 3xTg-AD mouse model of Alzheimer’s disease. Behav Brain Res 289:29–38.

Swerdlow RH (2007) Is aging part of Alzheimer’s disease, or is Alzheimer’s disease part of aging? Neurobiol Aging 28:1465–1480.

Tomás Pereira I, Gallagher M, Rapp PR (2015) Head west or left, east or right: interactions between memory systems in neurocognitive aging. Neurobiol Aging 36:3067–3078.

Villarreal JS, Dykes JR, Barea-Rodriguez EJ (2004) Fischer 344 rats display age-related memory deficits in trace fear conditioning. Behav Neurosci 118:1166–1175.

Wang W-C, Dew ITZ, Cabeza R (2015) Age-related differences in medial temporal lobe involvement during conceptual fluency. Brain Res 1612:48–58.

Widman AJ, McMahon LL (2018) Disinhibition of CA1 pyramidal cells by low-dose ketamine and other antagonists with rapid antidepressant efficacy. Proc Natl Acad Sci USA 115:E3007–E3016.

Wiltgen BJ, Sanders MJ, Anagnostaras SG, Sage JR, Fanselow MS (2006) Context fear learning in the absence of the hippocampus. J Neurosci 26:5484–5491.

Wright CI, Dickerson BC, Feczko E, Negeira A, Williams D (2007) A functional magnetic resonance imaging study of amygdala responses to human faces in aging and mild Alzheimer’s disease. Biol Psychiatry 62:1388–1395.

Yaffe K, Vittinghoff E, Lindquist K, Barnes D, Covinsky KE, Neylan T, Kluse M, Marmar C (2010) Posttraumatic stress disorder and risk of dementia among US veterans. Arch Gen Psychiatry 67:608–613.

Yehuda R, Golier JA, Tischler L, Stavitsky K, Harvey PD (2005) Learning and memory in aging combat veterans with PTSD. J Clin Exp Neuropsychol 27:504–515.

Zeng Y, Liu Y, Wu M, Liu J, Hu Q (2012) Activation of TrkB by 7,8-dihydroxyflavone prevents fear memory defects and facilitates amygdalar synaptic plasticity in aging. J Alzheimers Dis 31:765–778.

Zhan J-Q, Zheng L-L, Chen H-B, Yu B, Wang W, Wang T, Ruan B, Pan B-X, Chen J-R, Li X-F, Wei B, Yang Y-J (2018) Hydrogen Sulfide Reverses Aging-Associated Amygdalar Synaptic Plasticity and Fear Memory Deficits in Rats. Front Neurosci 12:390.

